# Caloric Restriction Remodels Energy Metabolic Pathways of Gut Microbiota and Promotes Host Autophagy

**DOI:** 10.1101/2020.08.16.251215

**Authors:** Yuping Yang, Shaoqiu Chen, Yumin Liu, Yuanlong Hou, Xie Xie, Xia Zhang, Aihua Zhao, Xiaojiao Zheng, Jiajian Liu, Tianlu Chen, Tianma Yuan, Hongjing Yu, Chongchong Wang, Yifan Sun, Jingcheng Wang, Xiaoyan Wang, Wei Jia

## Abstract

Calorie restriction (CR) can improve the metabolic balance of adults and elevate the relative abundance of probiotic bacteria in the gut while promoting longevity. However, the interaction between remodeled intestinal flora and metabolic improvement, as well as the mechanism for probiotic bacterial increase, are still unclear. In this study, using a metabolomics platform, we demonstrate for the first time, that CR leads to increased levels of malate and its related metabolites in biological samples. Next, we investigated the effects of CR on the gut microbial genome and the expression of mRNA related to energy metabolism which revealed a partially elevated TCA cycle and a subsequently promoted glyoxylate cycle, from which large amounts of malate can be produced to further impact malate related pathways in the host liver. Through the identification of key “hungry” metabolites produced by the gut microbiota that function in the promotion of autophagy in the host, further insight has been gained about a functional metabolic network important for both host-microbial symbiosis and maintenance of host health.

## Introduction

Caloric restriction (CR) refers to a 10-40% reduction in energy intake without nutritional deficiency^1–3^. It is known to prevent or reverse obesity to improve health, and is also one of the most common strategies for prolonging lifespan ^4,5 6^. CR counteracts the effects of over nutrition with an altered metabolism that is now considered a hallmark. Even though the mechanism by which CR works remains controversial, the cause is simply the reduction of total diet amount. CR plays a critical role in aging retardation by inducing metabolic reprogramming that involves multiple pathways and enzymes ^7,8^. Additionally, the microbiome impacts strongly on host metabolic phenotypes, because gut microbiota are associated with essential various biological functions and the microbial-host co-metabolism is necessary to process nutrients^9^. Long term CR extends lifespan and increases the abundance of gut probiotic bacteria, contributing to the health-benefiting effects of CR. ^10^ However, the reason for the increased proliferation of probiotic species in the gut under caloric restriction is still unclear. Given the importance of the metabolic environment on microbial growth, understanding the interaction between metabolism and flora might lead to a solution for enhancing the growth of beneficial bacteria in the gut.

Nutrition and energy intake status significantly affects the structures and functions of the gut microbial community.^11^ Certain bacteria have been reported to restrain multiple biosynthetic pathways and upregulate tricarboxylic acid (TCA) cycle genes in mouse under carbon starvation conditions.^12^ However, in addition to the TCA cycle, the microbial metabolism system can utilize another energy production pathway, the glyoxylate cycle (glyoxylate shunt), which bypasses carbon dioxide generation catalyzed by isocitrate and α-ketoglutarate dehydrogenase enzymes in the TCA cycle and thus conserves carbon atoms for gluconeogenesis. ^13,14^ The glyoxylate cycle is a metabolic function of both facultative anaerobic and aerobic bacteria and is essential for the production of bacterial acetate and fatty acid metabolism^13^. We hypothesized that when gut bacteria were under CR stress, they would take advantage of the glyoxylate cycle to reduce energy consumption and promote energy production. On the other hand, the series of metabolic changes caused by CR may be responsible for the increased abundance of probiotic bacteria which enhance physiological functions beneficial to host health to achieve a prolonged life.

We use metabolomic technology combined with high-throughput gut bacteria metagenomic and metatranscriptomic sequencing platforms to find variations in energy metabolism related pathways, involving the glyoxylate cycle and TCA cycle metabolisms induced by CR in experimental animals. By verifying the action of key “hungry” metabolites in the reconstruction of the gut microbiota community, we were able to identify certain interactions between specific metabolites and intestinal bacteria that defined a functional metabolic mechanism which contributes to host-microbial symbiosis and maintenance of host health.

## Results

### Metabolome and key metabolites variation Induced by 60% caloric restriction

60w-CR (CR for 60 weeks) and 40d-CR (CR for 40 days) calorie restriction (CR: 60% quantity of normal diet, the diet composition listed in Table.S1) were administered individually to two batches of Wistar rats. During 60 weeks of CR, concentrations of 32 metabolites in the urine of CR rats significantly differed from those of their normal diet controls (C) (Fig.1A). The largest FCs (fold changes of CR/C) were detected in gut microbiota metabolites, energy metabolism-related metabolites, and amino acids suggesting that CR had significant effects on host and gut microbiota energy metabolisms. Some TCA intermediates, malate, alpha-ketoglutarate and fumarate, peaked at the W60 time-point and malate had the most significant rise. However, the TCA cycle starting substrate, citrate and its metabolite, isocitrate were markedly decreased after W10 relative to controls. The observed increase in amino acids also suggested that CR induced tissue catabolism. Several amino acids, such as aspartate, valine, leucine and isoleucine showed significant variation after CR. Degradation products of these amino acids produce substrates for the TCA cycle.^15^. The urinary bacterial metabolites with increased FCs were, trans-ferulate, oxalate, 4-hydroxycinnamate and indole-3-acetate.

Malate, an energy metabolite and an important TCA intermediate, was also detected at higher concentrations in serum, liver and intestine of CR rats relative to controls at both Week 60 and Day 40(time of animal sacrifice). A separate, shorter time course experiment was performed (40 days) which confirmed the results of the long-term CR urine profile results (60 w) (Fig.1B). During the first four weeks of CR rats in the long-term CR experiment, dietary intake gradually decreased from 100% to 60% (From Week 0 to Week 4 with a decrease rate of 10%/wk) and malate levels increased significantly from the third week (dietary restriction 70%), and reached a very significant level in the fourth week (Fold change 5.5, *p* < 0.01) (Fig.S1 A). Urine malate levels at 3 weeks and 8 weeks after diet recovery at the 52th week were compared with C group and the results showed that the malate levels of CR-R rats decreased and returned to normal after a 3 and 8 week period of diet recovery (Fig.S1B). Measured malate levels of serum, ileum and colon in the CR-R group was lower than the CR group and closer to those of the C group after a 24 week period of diet recovery (Fig.1C). The elevated malate level observed in CR rats was lowered significantly both in liver and colon contents by vancomycin administration (antibiotics) (Fig.1D) indicating that bacterial metabolism was responsible for the malate production. The concentrations of succinate and aspartate were all increased in serum, ileum and colon contents of the CR rats (Fig.S1C).

Meanwhile, significant lower malate levels (p<0.05) were detected in both ileum and colon of CR (for 26 weeks) C57BL/6 mice compared with normal diet mice (Fig.S2).

**Fig.1.**
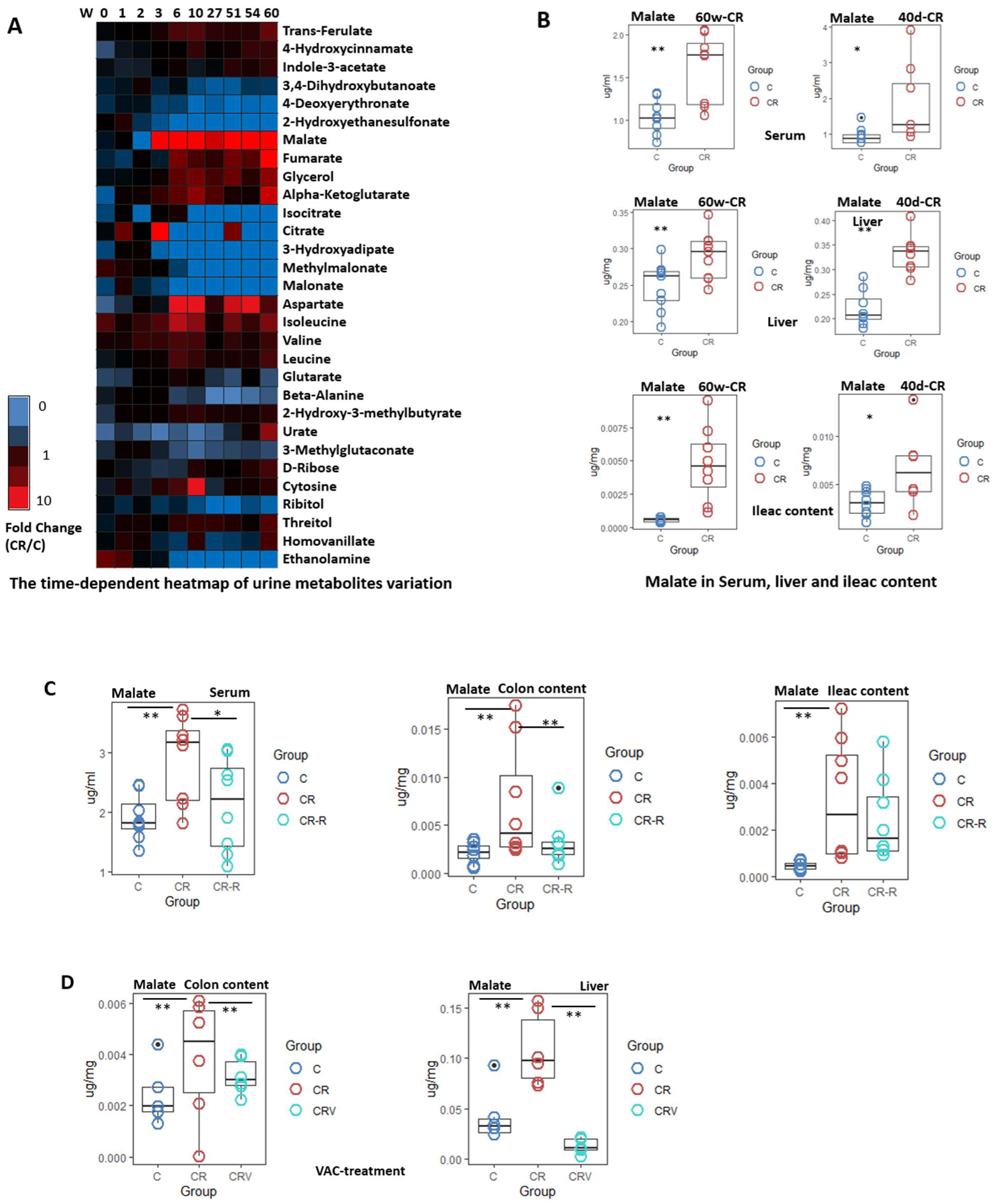
Key metabolite variations in rats of 60% calorie restriction (CR) A) The time-dependent heatmap of urine metabolite variation during long-term CR; B) Malate levels in serum, liver and ileum after long-term and short-term CR; C) Tissue malate level changes after 24 week diet recovery; D) Malate variances of Antibiotics (vancomycin administration for 5 days) intervention on CR experiments; *, T test (p<0.05),**,T test(p<0.01)

### Gut microbiota taxonomic changes induced by calorie restriction in rats

For 16S rRNA gene sequencing, ileum contents were collected at the end of the 60w-CR experiment and gut microbiota composition was analyzed by sequencing the V4 region of the 16S rRNA gene. Using 97% as a homology cut-off value, we obtained an averageof 17,329 reads per sample and 9,820 species-level operational taxonomic units (OTUs). The C group had 49231 OTUs, and the CR group had 42894 OTUs, representing a ~13% decrease (Fig.2A). The characteristic changes observed in the 16S rRNA gene sequencing results of the ileum flora induced by 60w-CR are displayed in Fig. S3. Microbiome taxonomic comparisons between C and CR groups at the phylum and genus level are shown (Fig. S3A and S3B, respectively). The main genera of gut bacteria increased in CR samples compared with C samples (CR/C (FC) > 1.2) and their relative abundances in CR samples are summarized in Table 1. The relative abundances of probiotic genera including, *Bifidobacterium*, *Prevotella*, *Parabacteroides*, *Lactobacillus* were elevated by CR.

**Fig. 2.**
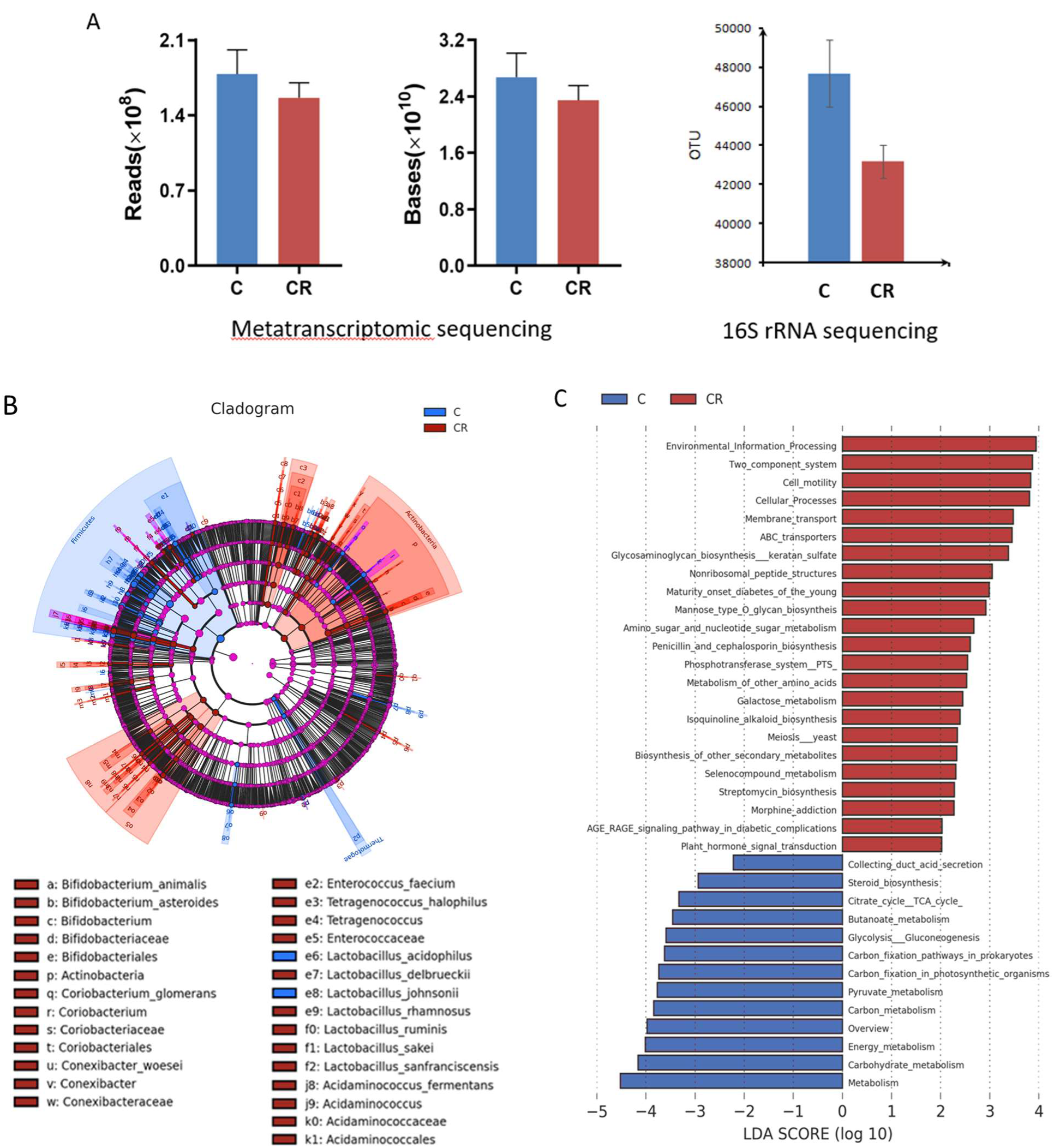
The characteristic changes of gut flora induced by calorie restriction. A)Total numbers of OUT comparison of C and CR groups by 16S rRNA sequencing and metatranscriptomic sequencing; B) LEfSe analysis of Microbiome structure and C) Functional group diagram between C and CR groups

**Tab. 1.** Predominantly increased gut bacteria (FC > 1.2) and their percentage of all abundance in CR samples of 60w-CR

| 16S rRNA sequencing |  |  |
| --- | --- | --- |
| Name | CR/C (FC) | % in CR group |
| g__Bifidobacterium | 12.27 | 0.21 |
| g__Prevotella | 10.76 | 0.70 |
| g__Parabacteroides | 7.05 | 0.06 |
| g__Lactobacillus | 6.93 | 23.71 |
| g__Marvinbryantia | 6.08 | 0.12 |
| g__Helicobacter | 5.43 | 0.18 |
| g__Allobaculum | 4.13 | 0.89 |
| g__Candidatus_Saccharimonas | 3.64 | 2.91 |
| g__Clostridium_sensu_stricto_1 | 3.26 | 4.57 |
| g__Desulfovibrio | 1.96 | 0.54 |
| g__Papillibacter | 1.53 | 0.02 |
| g__Ruminococcus | 1.47 | 1.54 |
| g__Bacteroides | 1.44 | 0.60 |
| g__Dorea | 1.33 | 0.37 |
| Metagenomic sequencing |  |  |
| Name | CR/C | % in CR group |
| Lactobacillus_murinus | 3.14 | 2.12 |
| Ruminococcus_bromii_L2-63 | 1.52 | 0.48 |
| Lactobacillus_gasseri | 1.64 | 0.42 |
| uncultured_prokaryote | 3.60 | 0.22 |
| uncultured_Clostridiales_bacterium | 1.37 | 0.19 |
| uncultured_Firmicutes_bacterium | 2.12 | 0.14 |

In the metagenomic sequencing protocol, the gene sequences obtained by high-throughput sequencing are optimized to obtain an average clean data set consisting of 2.0E+08 reads and 3.0 E+10 bases. The sequence construction database of bacteria, fungi, archaea and viruses was screened using the NT database of NCBI, the UniSeq sequence was compared with the constructed microbial database and the comparison results were then imported into Megan software to obtain the species classification of each sequence (differences are shown in Figs. S4A,B,C). The strains with different expressions (CR/C (FC) > 1.2) between the two groups are listed in Table 2.

**Table 2.** Corresponding gene labelling in Fig.4A from Metatranscriptomic sequencing

|  | KO_L3 | Definition | name.EC | Average C | Average CR | FC<sub>(CR/R)</sub> | P value |
| --- | --- | --- | --- | --- | --- | --- | --- |
| ① | Citrate cycle (TCA cycle)<br>[PATH:ko00020] | [EC:4.1.1.49]<br>pckA | phosphoenolpyruvate carboxykinase (ATP) | 408146.41 | 284877.75 | 0.70 | 0.14 |
|  | Citrate cycle (TCA cycle)<br>[PATH:ko00020] | [EC:4.1.1.32]<br>pckA, PCK | phosphoenolpyruvate carboxykinase (GTP) | 6322.31 | 5557.02 | 0.88 | 0.65 |
| ② | 00250 Alanine, aspartate and<br>glutamate metabolism<br>[PATH:ko00250] | [EC:2.6.1.1]<br>GOT1 | aspartate aminotransferase, cytoplasmic | 16.77 | 67.06 | 4.00 | 0.17 |
|  | 00250 Alanine, aspartate and<br>glutamate metabolism<br>[PATH:ko00250] | [EC:2.6.1.1]<br>aspB | aspartate aminotransferase | 160.11 | 200.83 | 1.25 | 0.42 |
| ③ | Citrate cycle (TCA cycle)<br>[PATH:ko00020] | [EC:6.4.1.1]<br>pycB | pyruvate carboxylase subunit B | 312.03 | 332.24 | 1.06 | 0.83 |
| ④ | Citrate cycle (TCA cycle)<br>[PATH:ko00020] | [EC:1.2.4.1]<br>PDHA, pdhA | pyruvate dehydrogenase E1 component alpha<br>subunit | 146.35 | 69.48 | 0.47 | 0.04 |
|  | Citrate cycle (TCA cycle)<br>[PATH:ko00020] | [EC:1.2.4.1]<br>PDHB, pdhB | pyruvate dehydrogenase E1 component beta<br>subunit | 1963.84 | 1089.29 | 0.55 | 0.13 |
|  | Citrate cycle (TCA cycle)<br>[PATH:ko00020] | [EC:2.3.1.12]<br>DLAT, aceF, pdhC | pyruvate dehydrogenase E2 component<br>(dihydrolipoamide acetyltransferase) | 122.39 | 107.94 | 0.88 | 0.82 |
|  | Citrate cycle (TCA cycle)<br>[PATH:ko00020] | [EC:1.2.7.1]<br>por, nifJ | pyruvate-ferredoxin/flavodoxin oxidoreductase | 856328.77 | 571794.50 | 0.67 | 0.12 |
| ⑤ | 00620 Pyruvate metabolism<br>[PATH:ko00620] | [EC:6.2.1.13] | acetyl-CoA synthetase (ADP-forming) | 108.01 | 50.17 | 0.46 | 0.55 |
| ⑥ | 00620 Pyruvate metabolism<br>[PATH:ko00620] | [EC:6.2.1.1]<br>ACSS, acs | acetyl-CoA synthetase | 37.70 | 19.90 | 0.53 | 0.52 |
| ⑦ | Glyoxylate and dicarboxylate<br>metabolism [PATH:ko00630] | [EC:2.3.3.1]<br>CS, gltA | citrate synthase | 114.96 | 179.53 | 1.56 | 0.69 |
| ⑧ | Glyoxylate and dicarboxylate<br>metabolism [PATH:ko00630] | [EC:4.2.1.3]<br>ACO, acnA | aconitate hydratase | 658.97 | 768.77 | 1.17 | 0.48 |
| ⑨ | Citrate cycle (TCA cycle)<br>[PATH:ko00020] | [EC:1.1.1.41]<br>IDH3 | isocitrate dehydrogenase (NAD<sup>+</sup>) | 36.92 | 22.56 | 0.61 | 0.42 |
|  | Citrate cycle (TCA cycle)<br>[PATH:ko00020] | [EC:1.1.1.42]<br>IDH1, IDH2, icd | isocitrate dehydrogenase | 67.97 | 30.80 | 0.45 | 0.22 |
| ⑩ | Citrate cycle (TCA cycle)<br>[PATH:ko00020] | [EC:1.2.7.3<br>1.2.7.11]<br>korA, oorA, oforA | 2-oxoglutarate/2-oxoacid ferredoxin<br>oxidoreductase subunit alpha | 1423.83 | 1301.89 | 0.91 | 0.77 |
|  | Citrate cycle (TCA cycle)<br>[PATH:ko00020] | [EC:1.2.7.3<br>1.2.7.11]<br>korB, oorB, oforB | 2-oxoglutarate/2-oxoacid ferredoxin<br>oxidoreductase subunit beta | 2635.85 | 2409.43 | 0.91 | 0.85 |
|  | Citrate cycle (TCA cycle)<br>[PATH:ko00020] | [EC:1.2.7.3]<br>korC, oorC | 2-oxoglutarate ferredoxin oxidoreductase<br>subunit gamma | 149.74 | 137.17 | 0.92 | 0.85 |
| ⑪ | Citrate cycle (TCA cycle)<br>[PATH:ko00020] | [EC:6.2.1.4<br>6.2.1.5]LSC2 | succinyl-CoA synthetase beta subunit | 253.70 | 475.66 | 1.87 | 0.33 |
|  | Citrate cycle (TCA cycle)<br>[PATH:ko00020] | [EC:6.2.1.5]<br>sucD | succinyl-CoA synthetase alpha subunit | 417.32 | 511.81 | 1.23 | 0.51 |
|  | Citrate cycle (TCA cycle)<br>[PATH:ko00020] | [EC:6.2.1.5]<br>sucC | succinyl-CoA synthetase beta subunit | 55.92 | 357.88 | 6.40 | 0.01 |
|  | Citrate cycle (TCA cycle)<br>[PATH:ko00020] | [EC:6.2.1.4 6.2.1.5]<br>LSC1 | succinyl-CoA synthetase alpha subunit | 282.46 | 461.40 | 1.63 | 0.43 |
| ⑫ | Citrate cycle (TCA cycle)<br>[PATH:ko00020] | [EC:1.3.5.1 1.3.5.4]<br>sdhB, frdB | succinate dehydrogenase / fumarate<br>reductase, iron-sulfur subunit | 2381.50 | 1999.44 | 0.84 | 0.58 |
|  | Citrate cycle (TCA cycle)<br>[PATH:ko00020] | [EC:1.3.5.1 1.3.5.4]<br>sdhA, frdA | succinate dehydrogenase / fumarate<br>reductase, flavoprotein subunit | 20.87 | 44.64 | 2.14 | 0.08 |
|  | Citrate cycle (TCA cycle)<br>[PATH:ko00020] | [EC:1.3.5.4]<br>frdA | fumarate reductase flavoprotein subunit | 8129.70 | 7911.62 | 0.97 | 0.89 |
|  | Citrate cycle (TCA cycle)<br>[PATH:ko00020] | [EC:1.3.5.4]<br>frdB | fumarate reductase iron-sulfur subunit | 3.81 | 0.00 | 0.00 | 0.35 |
| ⑬ | Citrate cycle (TCA cycle)<br>[PATH:ko00020] | [EC:4.2.1.2]<br>E4.2.1.2AA, fumA | fumarate hydratase subunit alpha | 1466.07 | 1936.33 | 1.32 | 0.20 |
|  | Citrate cycle (TCA cycle)<br>[PATH:ko00020] | [EC:4.2.1.2]<br>E4.2.1.2AB, fumB | fumarate hydratase subunit beta | 24.87 | 105.70 | 4.25 | 0.00 |
|  | Citrate cycle (TCA cycle)<br>[PATH:ko00020] | [EC:4.2.1.2]<br>E4.2.1.2B, fumC | fumarate hydratase, class II | 2558.91 | 1275.48 | 0.50 | 0.08 |
| ⑭ | Citrate cycle (TCA cycle)<br>[PATH:ko00020] | [EC:1.1.1.37]<br>MDH1 | malate dehydrogenase | 1481.68 | 1836.30 | 1.24 | 0.71 |
|  | Citrate cycle (TCA cycle)<br>[PATH:ko00020] | [EC:1.1.1.37]<br>mdh | malate dehydrogenase | 535.21 | 156.04 | 0.29 | 0.24 |
| ⑮ | Glyoxylate and dicarboxylate<br>metabolism [PATH:ko00630] | [EC:3.1.3.18<br>3.1.3.48]<br>PGP, PGLP | phosphoglycolate phosphatase | 16.45 | 83.12 | 5.05 | 0.03 |
|  | Glyoxylate and dicarboxylate<br>metabolism [PATH:ko00630] | [EC:3.1.3.18]<br>gph | phosphoglycolate phosphatase | 2966.02 | 2367.72 | 0.80 | 0.43 |
| ⑯ | Glyoxylate and dicarboxylate<br>metabolism [PATH:ko00630] | [EC:1.1.3.15]<br>glcD | glycolate oxidase | 226.18 | 593.51 | 2.62 | 0.02 |
| ⑰ | 00620 Pyruvate metabolism<br>[PATH:ko00620] | [EC:1.1.1.40]<br>maeB | malate dehydrogenase (oxaloacetate-<br>decarboxylating) (NADP+) | 173.49 | 253.63 | 1.46 | 0.08 |

The metatranscriptomic sequencing results revealed that gut contents from CR rats had a higher total number of reads and total number of bases (Fig.2A), which were consistent with the above 16S rRNA gene sequencing results. The differences of flora species group diagram with significant differences between the two groups from LEfSe analysis are displayed in Fig.2B. The gut bacteria in CR samples most increased compared with C samples (CR/C (FC) > 1.2) and their relative abundances in CR samples are also summarized in Table 2. The relative abundances of probiotic genera including *Lactobacillus*, *Bifidobacterium* and *Parabacteroides* were elevated by CR.

### Gut microbiota functional transcriptomic changes induced by calorie restriction

At KEGG level 4 (gene expression level), a total of 214 KOs with significant differences in expression levels were enriched. Metatranscriptomic results indicated that metabolism pathways especially those related to energy metabolism, were inhibited in the gut flora of CR rats (Fig.2C). These inhibited pathways included carbohydrate metabolism, energy metabolism, carbon metabolism, pyruvate metabolism, carbon fixation pathways, glycolysis, butanoate metabolism and the TCA cycle (Fig S5 A). There were some pathways that were up-regulated (Fig S5 B) in CR rats. The functional annotation results of each Unigene in the KEGG database, combined with its corresponding expression, were used to analyze the expression distribution of samples in a given KEGG orthology (KO) level. Carbohydrate and amino acid metabolic pathways changed significantly, including pyruvate, dicarboxylate, alanine, aspartate, glutamate and butanoate metabolisms as well as the TCA and glyoxylate cycles (Table 2).

In the gluconeogenesis pathway, oxaloacetate is transformed into phosphoenolpyruvate and carbon dioxide via catalysis by phosphoenolpyruvate carboxylation kinase. Phosphoenolpyruvate carboxykinase (ATP) [EC:4.1.1.49] and (GTP) [EC:4.1.1.32] ① were found to be slightly decreased in the CR group. Pyruvate carboxylase catalyzes the reversible carboxylation of pyruvate to form oxaloacetate. Transcriptional expression of pyruvate carboxylase subunit B [EC:6.4.1.1] ③ was not changed by CR. The complex enzyme [EC:1.2.4.1] ④, which catalyzes the irreversible reaction of pyruvate to Acetyl-coA, was reduced by one-half in the CR group. 2-oxoglutarate / 2-oxoacid feredoxin oxidoreductase subunit alpha [EC: 1.2.7.3 1.2.7.11] ⑩, responsible for the biotransformation of pyruvate to Acetyl-coA in the CR group also did not exhibit significant changes. These results indicated that the pathway from pyruvate to Acetyl-CoA was inhibited. Transcriptional expression of acetyl CoA synthetase [6.2.1.13] ⑤ and [6.2.1.1] ⑥ was decreased in CR group.

In TCA cycle pathways, the transcription expression of aconitate hydratase [EC: 4.2.1.3] ⑧and citrate synthase [EC: 2.3.3.1] ⑦ were higher in gut bacteria of the CR group relative to the C group, indicating that the metabolic pathway from isocitrate to oxaloacetic acid was activated by calorie restriction. On the other side of the cycle, the levels of succinyl CoA synthetase, Alpha & beta subunits [EC: 6.2.1.4&6.2.1.5] ⑪ were all up-regulated, consistent with the results of the metagenome. Fumarate formation, catalyzed by succinate dehydrogenase / fumarate reductase, flavoprotein subunit [EC: 1.3.5.1&1.3.5.4] ⑫, was not significantly elevated. Next, fumarate hydratase Alpha & beta [EC: 4.2.1.2] ⑬ which catalyzes fumarate to malate, was not significantly increased, consistent with the metagenome results. Malate dehydrogenase [EC: 1.1.1.37] ⑭ did not significantly increase. At the other end of the TCA cycle, isocitrate dehydrogenase [EC: 1.1.1.42] and isocitrate dehydrogenase (NAD +) [EC: 1.1.1.41] ⑨ were both decreased in CR group, indicating that the biotransformation of isocitrate to α – ketoglutarate was inhibited, consistent with the result of metagenome.

The results of the metatranscriptome indicated that gene expression levels of compounds in the glyoxylate pathway, another malate related metabolic pathway, had changed in intestinal microorganisms. Aspartate aminotransferase, cytoplasmic [EC: 2.6.1.1] ②, a key enzyme that catalyzes the biotransformation of aspartate into oxaloacetate which in turn, was transformed into citrate and was increased in expression corresponding to the observed levels of aspartate and citrate in the CR group. Meanwhile, two irreversible enzymes, phosphoglycolate phosphotase [EC: 3.1.3.18] ⑮ and glycolate oxidase [EC:1.1.3.15] ⑯, which catalyze the transformation of phosphoglycolate to glyoxylate, were significantly up-regulated, indicating caloric restriction significantly elevated the production of glyoxylate. Malic enzyme 1[EC:1.1.1.40] ⑰ which was elevated in gut bacteria, catalyzes the reversible oxidative decarboxylation of malate to pyruvate(Fig.3A).

**Fig. 3.**
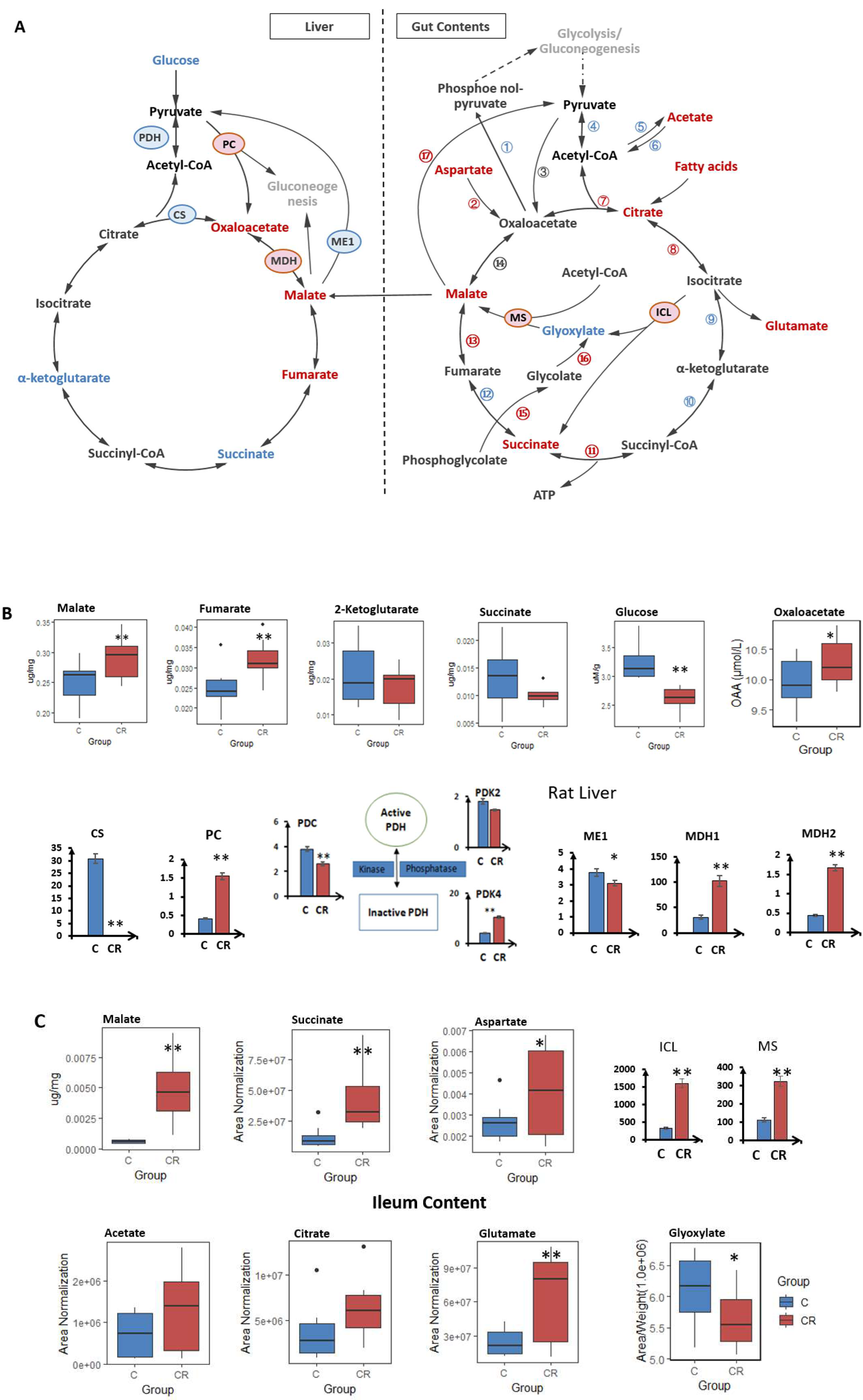
Malate-related enzymes and metabolite changes in Host hepatic and ileum contents induced by calorie restriction. A) malate-related pathways both in liver and ileum contents, metabolic analysis integrated with Metatranscriptomic functional annotation (Circled numbers listed in Table 2) to construct the TCA and glyoxylate cycle pathways(Red: up-regulated in CR group; Blue: down-regulated in CR group); B) Boxplots of malate-related enzymes and metabolites in rat livers; C) Boxplots of malate-related enzymes and metabolites in rat ileum contents; MS: Malate synthase; ICL: Isocitrate lyase; MDH1: Malate dehydrogenase 1; MDH2: Malate dehydrogenase 2; ME1: Malic enzyme 1; PC: Pyruvate carboxylase; CS: Citrate synthase; PDC: Pyruvate dehydrogenase complex; PDK2: Pyruvate dehydrogenase kinase isozyme 2; PDK4: Pyruvate dehydrogenase kinase isozyme 4 (CR compared with C group: *, T test (p<0.05); **, T test (p<0.01)

Functional annotation of the metagenome was constructed using the KEGG biological pathway database. Differentially expressed functional genes related to the TCA and glyoxylate cycle pathways are listed in Fig. S4D. Some other differentially expressed transcription genes detected by metatranscriptomic sequencing are listed in Table S2.

### Variations of key metabolites, gene and transcription expressions of key enzymes were detected in the liver and ileum contents of 60w-CR rats

In mammals, the liver is the center for malate metabolism and the TCA cycle (Fig.3A). The mRNA expression levels of two malate dehydrogenases (MDH1 and MDH2) were 3.32 and 3.80 fold higher, respectively, in the livers of the CR rats relative to controls (Fig.3B). Pyruvate is a downstream metabolite of the malate dehydrogenases, and the mRNA levels of pyruvate carboxylase (PC) and pyruvate dehydrogenase kinase isozyme 4 (PDK4) were also increased 3.74 and 2.60 fold, respectively, in the livers of CR rats compared to controls. Conversely, the mRNA levels of citrate synthase (Cs) and pyruvate dehydrogenase complex (PDC) decreased to 0.3% and 69% of control values, respectively, in the CR rat liver. Hepatic glucose was also decreased in CR rats. Malate concentrations were consistently found to be elevated. Fumarate, a precursor to malate in the TCA cycle, was increased by CR while the concentrations of succinate and 2-ketoglutarate decreased to 0.78 fold and 0.83 fold, respectively. Oxaloacetate levels increased to 1.5 fold in CR rat livers compared to the C group.

Metabolomic analysis showed that 2 key metabolites in the glyoxylate cycle, malate (+677%) and citrate (+166%) increased in the ileum content of the CR rats with a corresponding 20% decrease in glyoxylate levels (Fig.3C). The concentrations of 4 metabolites that are relevant to the glyoxylate cycle, aspartate (+153%), acetate (+188%), glutamate (+264%), and succinate (+294%) were also increased in the ileum content of the CR rats. The concentrations of acetate (+193%), succinate (+332%), aspartate (+472%), and glutamate (+226%) were increased in the colon content of the CR rats (Fig S1C). Concentrations of short chain fatty acids, such as propanoate (+278%), butyrate (+195%), and pentanoate (+252%) were increased in the feces of the CR rats (Fig.S6). However, oxaloacetate could not be detected in any gut contents. In order to validate the outcome of metagenomics, qPCR detection was used to quantify the gene expression levels of key enzymes involved in the glyoxylate cycle. The DNA expression levels of isocitratelyase (ICL) and malate synthase (MS) were elevated by 4.81 fold (p=0.005) and 2.86 fold (p=0.009), respectively.

### Metabolomic changes caused by acetate stimulation in gut contents cultures

The presence of acetic acid was considered an important condition for activation of the glyoxylate cycle.^13^ Further, in order to verify that glyoxylate activation in intestinal flora was the cause of malic acid increase, acetic acid was added to the gut contents cultures derived from rats in groups C and CR. The survival of bacteria stimulated by acetic acid and any accompanying metabolic variations were then observed. The cultures treated with acetic acid were cultured for 1, 2, 3 hours and the conditioned media was collected for colony analysis at each time point. The metabolic analysis results showed that malate and fumarate levels were elevated and that the glyoxylate and succinate levels were decreased following acetate addition. Additionally, isoleucine and GABA were up-regulated, indicating that some amino acids were *de novo* synthesized in CR conditions (Fig.4A).

**Fig.4.**
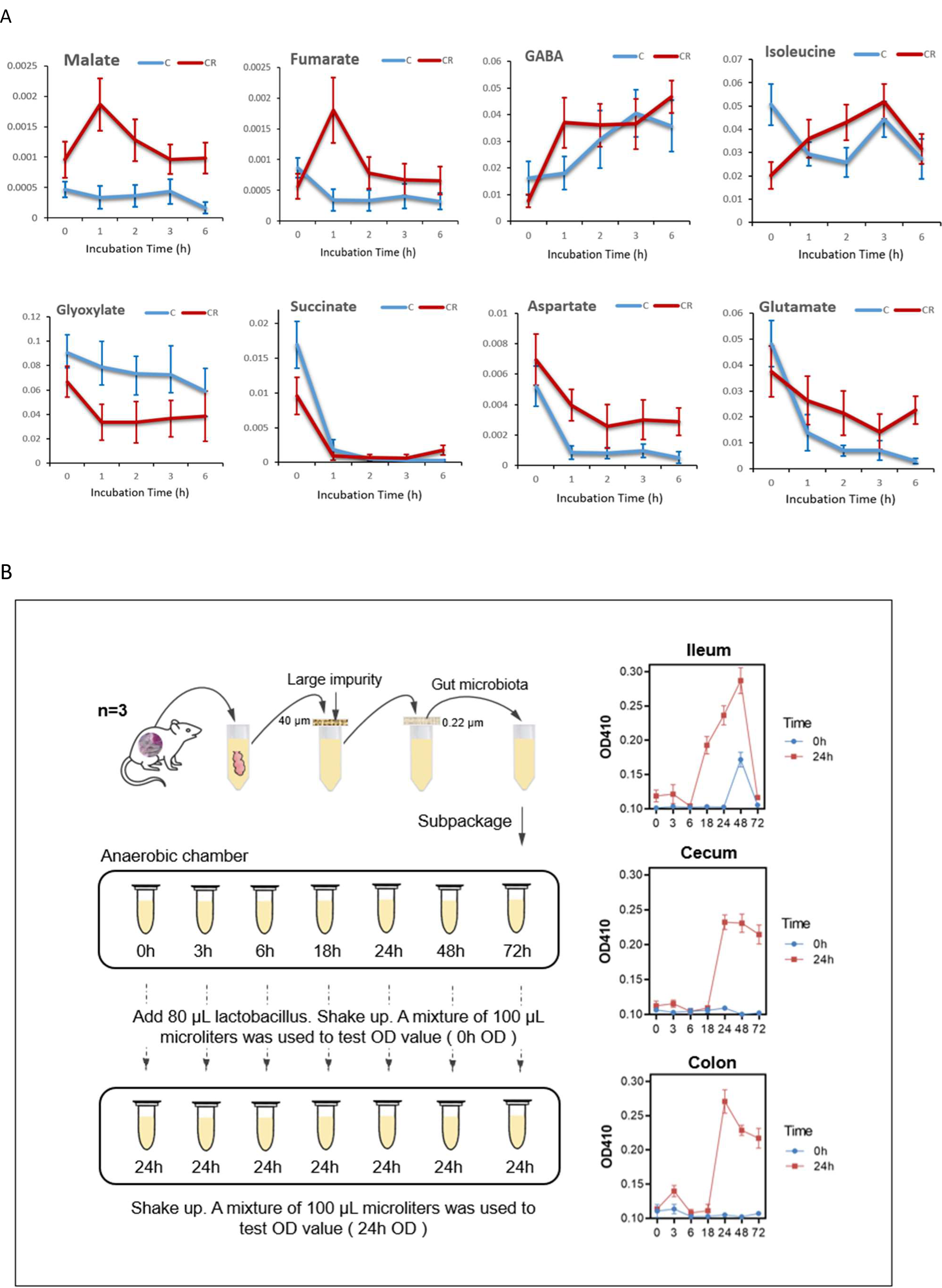
In vitro culture of gut microbiota: Metabolic changes culture medium *in vitro* stimulated by acetate (A) and the starvation cultured gut bacteria metabolites on *Lactobacillus murinus* proliferation (B).

### Starvation culture of the gut microbiota promotes the growth of Lactobacillus

In CR treated rats, a significantly increased abundance of *Lactobacillus* was found in several gut contents. The results of metabolomics revealed that Lactobacillus and other bacteria were positively correlated with malate (Table 3). After being isolated and cultured *in vitro*, *Lactobacillus murinus* was identified and cultured *in vitro*.

**Table 3.** Correlations between 16S rRNA sequencing ileum microbiota and malate level

| Taxonomical classification(Family) |  | Genus | Coefficient | Which malate level<br>Correlated with ileum<br>microbiota |
| --- | --- | --- | --- | --- |
| Order | Family |  |  |  |
| **Lactobacillales** | Lactobacillaceae | Lactobacillus | 0.79 | Colon |
| **Clostridiales** | Ruminococcaceae | Non | 0.66 | Colon |
| **Erysipelotrichales** | Erysipelotrichaceae | Allobaculum | 0.69 | Colon |
| **Clostridiales** | Non | Non | 0.76 | Ileum |
| **Unknown_Order** | Unknown_Family | Candidatus_Saccharimonas | 0.76 | Ileum |
| **Unknown_Order** | Unknown_Family | Candidatus_Saccharimonas | 0.75 | Ileum |
| **Clostridiales** | Ruminococcaceae | Non | 0.73 | Ileum |
| **Clostridiales** | Lachnospiraceae | Blautia | 0.70 | Ileum |
| **Clostridiales** | Lachnospiraceae | Blautia | 0.69 | Ileum |
| **Coriobacteriales** | Coriobacteriaceae | Adlercreutzia | 0.77 | Serum |
| **Lactobacillales** | Lactobacillaceae | Lactobacillus | 0.75 | Serum |
| **Erysipelotrichales** | Erysipelotrichaceae | Allobaculum | 0.75 | Serum |
| **Lactobacillales** | Lactobacillaceae | Lactobacillus | 0.75 | Serum |
| **Clostridiales** | Clostridiaceae_1 | Non | 0.73 | Serum |
| **Erysipelotrichales** | Erysipelotrichaceae | Allobaculum | 0.73 | Serum |
| **Clostridiales** | Clostridiaceae | Clostridium_sensu_stricto_1 | 0.72 | Serum |
| **Clostridiales** | Ruminococcaceae | Non | 0.69 | Serum |
| **Clostridiales** | Clostridiaceae | Clostridium_sensu_stricto_1 | 0.69 | Serum |

Contents of different intestinal segments from normal rats were harvested and cultured for 72 hours under strict anaerobic condition. At different time-points, the conditioned culture medium was removed and was used to culture *Lactobacillus murinus* isolated from CR rats. The *Lactobacillus* growth was assessed using UV absorption to measure optical density (OD) values (Fig.4B). The results showed that all media from the three gut segments obtained at starvation stage (18 hours later) had a higher absorbance, with a more than 2.5 fold change, indicating that starvation culture media promoted the proliferation of *Lactobacillus murinus*.

### Expression of autophagy- and mitophagy-related proteins during CR and sodium malate treatment

CR’s ability to stimulate autophagy and mitophagy was confirmed in CR rat hepatic tissues. The western blot results show that both LC3-I and LC3-II were significantly increased in CR rats (Fig.5). Specifically, pAMPK and p62 were all up-regulated in CR rat livers, indicating that CR was not only able to increase levels of pAMPK, one of the kinases involved in autophagy initiation, but also increased the accumulation of p62, an important component for the sequestration phase towards the formation of the final autophagosome. CR had no impact on autophagy related protein-7 (ATG7) (Fig.5A) levels. P62 generally changes in two ways. One is p62 can be degraded by autophagy, so p62 protein expression reflects the strength of autophagy. That is, when LC3-II is elevated and p62 decreases at the same time, autophagy is unobstructed. The other way is, LC3-II is elevated along with p62, indicating that autophagy starts normally but that events downstream are blocked due to lack of lysosome fusion.

The pAMPK/AMPK ratio was greatly increased in the CR group. Malate increased LC3-II expression in the human colon carcinoma, HCT116 cell line, and sodium malate had more effects than dimethyl malate (Fig. S7). Sodium malate increased both LC3-I and LC3-II in the human NCM460 cell line (Normal colon epithelial cells) (Fig.5B, C). Furthermore we found that both NCM460 cells and CR primary rat hepatocytes treated with sodium malate showed a dose-dependent elevation of the biomarker for autophagy and mitophagy (the ratio of LC3-II/β-ACTIN in the total cell lysate)^16^. These results showed that CR can induce and promote autophagy and that malate was able to promote autophagy and mitophagy *in vitro*.

**Fig. 5.**
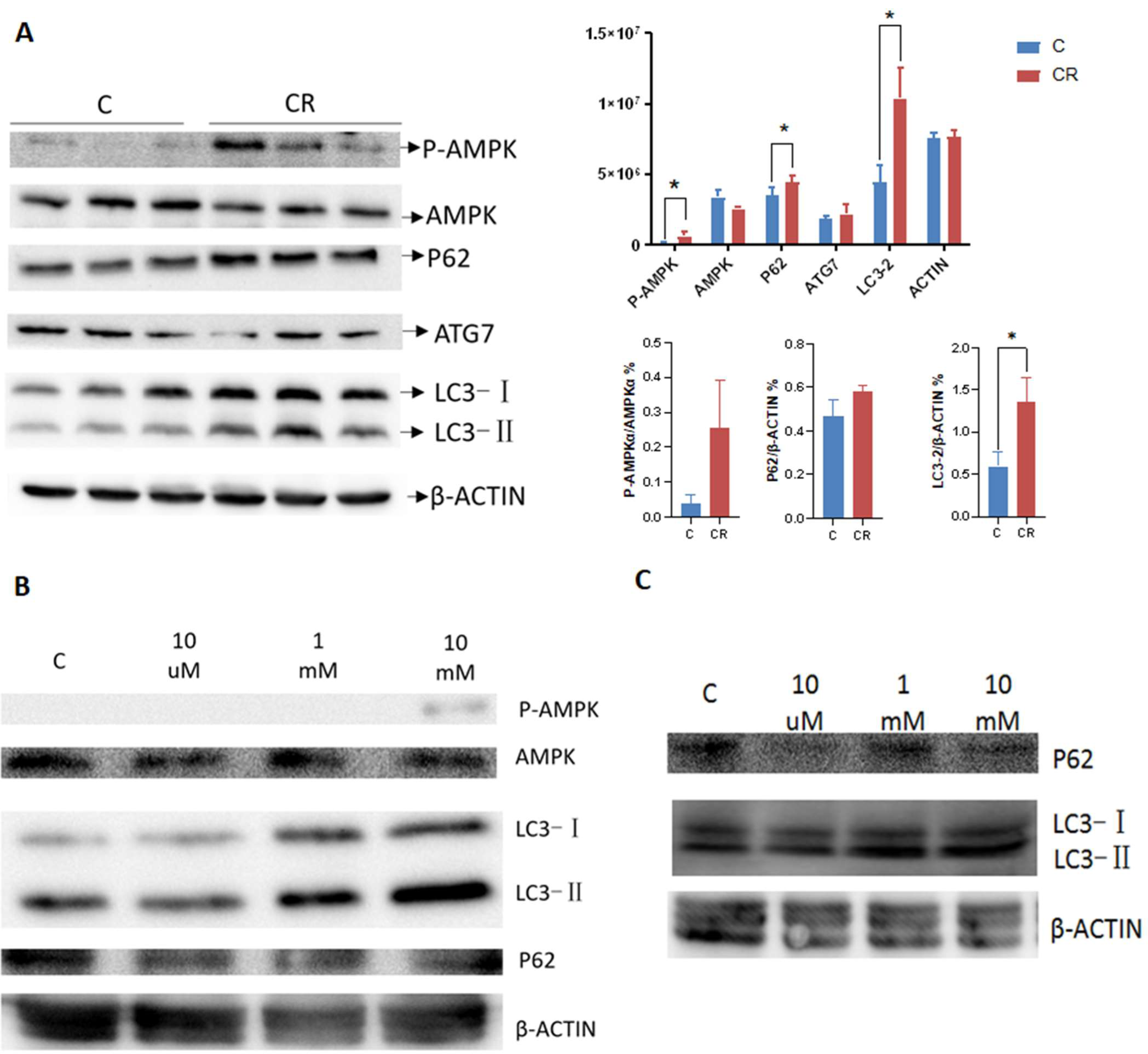
Autophagy and mitophagy induced by CR and malate in liver and cell line. A) Western blot of autophagy and mitophagy related proteins in liver tissues CR rats and controls (n = 3 biological replicates). B) Western blot of autophagy and mitophagy related proteins induced by sodium malate of different concentration in NCM460 cell lines. C) Western blot of p62 and LC3-II proteins induced by sodium malate of different concentration in primary rat hepatocytes.

## Discussion

How calorie restriction extends lifespan is not clear. The inverse correlation between food intake and the length of lifespan of mammals suggests an important role of energy metabolic regulation ^8^. Use of systematic screening tools, including analysis of the metabolome, microbiome and metagenome, proved to be effective in identifying changes that occur in metabolic pathways during CR. Although CR decreased the basal metabolic rate in our experiments, CR induced increased concentrations of malate along with a similar concentration trend for aspartate indicated that the important TCA cycle intermediate, oxaloacetate could continue to be produced at a relatively high rate. CR also increased malate levels in serum, urine, liver and gut contents of rats with similar results seen in human volunteers with calorie-restricted diet^17^. Malate has critical roles in both aerobic and anaerobic metabolism. Malate supplementation extended the lifespan of *Caenorhabditis elegans*^18^, and increased mitochondrial respiration and ATP production in human skeletal muscles^19^. Malate, oxaloacetate and aspartate also participate in the malate-aspartate shuttle to maximize ATP production from glycolysis, and to activate downstream targets of CR in yeast^20^. Additionally, aspartate levels were significantly elevated in the urine and ileum of CR rats. Both malate and aspartate are key metabolites in gluconeogenesis, which increases during CR^21^. Therefore, the higher levels of malate in CR rats may improve the efficiency of energy production, and compensate for the shortage of energy intake. Regarding the other metabolites in the hepatic TCA cycle, fumarate levels were increased but 2-ketoglutarate isocitrate and citrate levels decreased in liver during CR indicating that CR has a complex effect on the TCA cycle (Fig.3B). It was reported that the activity of malate dehydrogenase(MDH) was found to be increased in the liver mitochondria animals ^22^. Herein we showed that CR upregulated gene expression of both MDH1 (cytosolic), MDH2 (mitochondrial) and oxaloacetate in rat livers, suggesting that malate-to-oxaloacetate conversion might have increased both inside and outside of mitochondria (Fig.3B). Malic enzyme1 (ME1) catalyzes the biotransformation of malate to pyruvate in the cytoplasm, thus connecting the TCA cycle and glycolysis pathways^23^. The down-regulated gene expression of ME1 indicates that CR might have the effect of inhibiting the biotransformation of malate-to-pyruvate. The pyruvate dehydrogenase complex (PDC) irreversibly catalyzes the conversion of pyruvate to Acetyl-CoA in mitochondria, a key step connecting glycolysis and the TCA cycle. PDC gene expression was down-regulated in the livers of CR rats. Furthermore, pyruvate dehydrogenase kinase isozymes (PDKs), particularly PDK4, inactivate PDC^24^, and CR upregulated PDK4 gene expression. Therefore downregulated gene expression and enzymatic inactivation of PDC may both inhibit the TCA cycle via decreasing the pyruvate-to-Acetyl-CoA conversion. Meanwhile, upregulation of pyruvate carboxylase (PC) expression indicated that the first step in the initiation of gluconeogenesis occurs in CR animals to keep the body's blood sugar at a normal level ^25^. Furthermore, pyruvate carboxylase also participates in anaplerosis in that it catalyzes the bioconversion of pyruvate to oxaloacetate and thus replenishes the aspartate-malate shuttle and generates reducing equivalents reported to promote liver antioxidant capacity, another benefit that CR brings.^26^ Subsequently, the oxaloacetate level was raised in host liver (Fig.3A, B) and serum of different rat CR experiments (Fig. S6). Our results showed that gluconeogenesis was up-regulated in long-term CR rats, consistent with the previous reports. ^27,28^ But the entry of oxaloacetate into the TCA cycle is inhibited by the drastic downregulation of gene expression of citrate synthase (Cs), a regulatory enzyme that catalyzes the reaction of acetyl coenzyme A with oxaloacetate to produce citrate (Fig.3A, B). Oxaloacetate is also considered as a potent inhibitor of Complex II, and thus elevated oxaloacetate levels may further distort the TCA cycle^29^. Oxaloacetate is involved in gluconeogenesis, urea cycle, amino acid synthesis, and fatty acid synthesis, sufficient amounts of oxaloacetate thus may contribute to the body's building material supply during period of food shortage. Overall, the CR-induced changes in hepatic gene expression revealed an increased oxaloacetate production via activating malate metabolism and inhibiting the TCA cycle.

Our metabolomics analysis results showed that most of the urine metabolites decreased in the CR group, but the concentrations of malate and acetate increased. When the exogenous intake of energy intake is insufficient, the common goal of host and gut bacteria community becomes the promotion of gluconeogenesis to ensure the necessary energy supply. Different from animals, some plants and microorganisms contain glyoxysomes, in which the glyoxylate cycle opens up a saving mode to produce more carbohydrates.^30–32^ Little is known about its role in the communication between gut microbiota and the host. Both metatranscriptomics and metagenomics sequencing results showed that genes involved in the glyoxylate cycle were upregulated during CR, including phosphoglycolate phosphatase and glycolate oxidase (Table 2, Fig.S4). Metatranscriptomic results showed that energy metabolism including TCA cycle and pyruvate metabolism were generally decreased in CR gut bacteria (Fig.3C). However metabolomics results showed that parts of the TCA cycle participants were up-regulated in CR samples, such as malate, citrate and succinate. To find out where the elevated malate comes from, further metatranscriptomics functional analysis of the genes in TCA and glyoxylate cycle pathways was performed (Fig 4A, Table 2). The levels of succinyl CoA synthetase Alpha & beta subunit [EC: 6.2.1.4 & 6.2.1.5] ⑪ were all up-regulated. Under the catalytic control of succinyl CoA synthetase, succinate CoA is gets converted into succinate and coenzyme A. This process generates ATP or GTP, which is the only reaction in the TCA cycle accompanied by the formation of high-energy phosphate bonds, a result completely consistent with those found in the analysis of the metagenome. However, ⑫⑬⑭⑨ (Fig.3A) were unchanged or decreased in CR group, suggesting that the TCA of intestinal flora is not all elevated. However, these gene expression changes in the TCA metabolic pathway were not able to fully explain why intestinal malate levels rose significantly in CR groups.

The metatranscriptomic results also indicated that some gene expression in the glyoxylate pathway, another malate related metabolic pathway, had changed in intestinal microorganisms. Glyoxylate cycle and TCA cycle have multiple shared metabolites and enzymes. The increased levels of aspartate aminotransferase [EC: 2.6.1.1] ②, a key enzyme to initiate the glyoxylic acid cycle *via* its’ ability to catalyze the conversion of aspartate into oxaloacetate which subsequently forms citrate was in agreement with the increased level of aspartate and citrate in the CR group. Meanwhile, two irreversible enzymes, phosphoglycolate phosphotase [EC: 3.1.3.18] ⑮ and glycolate oxidase [EC:1.1.3.15] ⑯, which catalyze the transformation of phosphoglycolate to glyoxylate, were significantly up-regulated, indicating caloric restriction significantly promoted the production of glyoxylate. But metabolomics results showed that glyoxylate levels were lower in the CR group than C. To figure out this contradiction, the gene expression of two key enzymes, ICL and MS were detected by qPCR Isocitrate lyase (ICL) catalyzes the formation of glyoxlate and succinate from isocitrate. Glyoxylate is then subsequently reacted with acetyl-CoA to produce malate, catalyzed by MS (malate synthase) ^30^. Thus, the two key enzymes of the glyoxylate cycle, are ICL, an enzyme that channels metabolites into the glyoxylate cycle, and MS which is responsible for malate generation^33^. Our study showed that the copy numbers of ICL and MS genes were both significantly increased in CR rats (Fig.3A, C), suggesting that CR favored the glyoxylate cycle and malate production in gut microbiota as confirmed by the significant decreases of malate in urine and gut in the CR rats after antibiotic treatment (Fig. 1D). Glyoxylate was also detected in the gut contents of control group, indicating that the glyoxylic acid cycle does exist in intestinal bacteria. It was reported that intermittent fasting changes the composition of gut flora and increases fermentation in the gut to produce the metabolites, acetic acid and lactic acid. ^34^ *Acidaminococcus*, a Gram-negative coccus that consumes amino acids and produces acetic and butyric acids, was promoted by calorie restriction (Fig 3B).^35^ Similarly, in the CR group, the abundance of *Acidaminococcus*, and the acetic acid producing genus *Coriobacterium* was increased.^36^ The production of excessive acetate produced during starvation enables more Acetyl CoA to enter into the glyoxylate cycle *via* the metabolic enzymes of intestinal bacteria. In this way, glyoxylate is consumed and more malic acid is produced.^34^ This reaction occurs in all intestinal segments. Malic acid in ileum is absorbed into the host to further participate in whole-body metabolic process. L-malate administration improved the activities of mitochondrial oxidoreductases on aged rats.^37^ Acetate is absorbed and metabolized by intestinal cells to Acetyl CoA and promotes lipid β oxidation.^38^ The *in vitro* gut content culture with acetate intervention validated acetate’s effect on activation of the glyoxylate cycle to produce malate. These results confirmed that the initiation of the glyoxylate cycle pathway corresponded to the relative abundance of certain bacteria and that both of these factors were regulated by dietary energy intake.

The gut malic enzyme 1(ME1) [EC:1.1.1.40] ⑰ catalyzes the reversible oxidative decarboxylation of malate to pyruvate (Fig.3A, Table 2). The elevated ME1 found in gut bacteria indicated that the linkage between the glycolytic and citric acid cycles become intensified by CR in gut microbiota. Further, the higher levels of malate in the host doesn’t accelerate, or even inhibit the TCA cycle, because the concentrations of citrate and Isocitrate, the two downstream metabolites of malate in the TCA cycle were decreased during CR. Therefore, malate generated by gut bacteria enters the host circulation as an efficient energy source where it may act to alleviate the host’s energy production load and also decrease the production of reactive oxygen species (ROS) that are by products of oxidative phosphorylation. This is a plausible explanation for the CR-associated oxidative stress attenuation^39,40^.

Autophagy is a cellular process that digests damaged cellular components to meet energy demand when extracellular nutrients are scarce^41^, it is also a self-cleaning process that removes senescent organelles and maintains cellular function^42^. Autophagy is enhanced by inhibiting IGF-1/IR signaling^43,44^, and the latter is activated by nutrients, which further activate mTOR signaling thus deactivating the pro-autophagic serine/threonine protein kinase ULK1 (ULK1) complex and activating autophagy inhibitory S6 kinase (S6K). CR has been shown to induce autophagy in different organs and tissues ^45^, possibly by increasing AMP/ATP ratios and activating AMPK which inhibits mTOR and activates the ULK1 complex^41^. In our study, CR upregulated AMPK phosphorylation and protein expression of autophagy marker LC3-II in the liver, implicating that long-term CR may enhance the autophagy process. The role of malate in the induction of autophagy was studied with *in vitro* models, and we found that sodium malate dose-dependently deregulated protein expression of LC3-II, suggesting malate might have an active role in autophagy regulation (Fig.5). However, sodium malate did not change AMPK phosphorylation, possibly due to a relatively short period of exposure (12 hours). Further studies are warranted to confirm the role of malate in autophagy, and to fully illustrate the molecular mechanism.

CR reduced the richness of the gut microbiome and simplified its’ composition (Fig.2A). However, the abundance of many genera including those known to act as probiotics, *Bifidobacterium, Lactobacillus, Saccharimonas, Prevotella, Dorea and Ruminococcus*, were amplified in the CR gut, which is generally considered to be an important reason for CR to extend animals’ life ^46^. So what directly regulates the change of the composition of gut flora? Metabolic composition may be an answer. In the gut microenvironment, close interactions exist between metabolites and gut microbes. It was reported that *Ruminococcus*, *Prevotella* and *Lactobacillus* are all bacteria that grow more easily under enhanced malate conditions and that *Lactobacillus* reverts malic acid into lactic acid.^47^ So the contents of different intestinal segments from normal rats were used to explore the impact of “hungry gut metabolites” on bacteria growth. The results showed that the contents of ileum, cecum and colon under starvation culture conditions all significantly promoted the proliferation of *Lactobacillus murinus* (Fig.4B), proving that the CR-metabolic composition might be a key factor that impacts and even regulates gut flora composition.

This study uncovered CR induced gut microbial taxonomic and functional variations and their roles in producing metabolic benefits to the host. CR activates the glyoxylate cycle, which in turn produces a large amount of malate that supplements the supply of energy available to the host and the regulation of autophagy which increases the survival capability of cells existing in low nutrient environments. CR also promotes the increase in abundance of host beneficial, probiotic bacteria

## Method

### Animal Handling and Sampling

This study was conducted in accordance with the Chinese national legislation and local guidelines on laboratory animal welfare. It was performed at the Centre of Laboratory Animals, Shanghai Jiao Tong University, Shanghai, P. R. China. All of the Wistar rats and C57BL/6 mice were purchased from the Shanghai Laboratory Animal Co. (Sippr-BK, Shanghai, China), housed individually in stainless steel wire mesh cages, and provided with certified standard rat chow and tap water *ad libitum*. Room temperature and humidity were regulated at 24 ± 1 °C and 45 ±15%, respectively. A 12 h on/off light cycle was used with lights coming on at 8 a.m.

For the 60w-CR study, after 4 weeks of acclimatization in metabolic cages, 36 six-month-old male Wistar rats were randomly divided into the following three groups: (1) Control group (C group), access to food *ad libitum* and (2) Calorie Restriction group (CR group). At the beginning of CR treatment, food intake was decreased to 90% of normal food intake in the first week, 80% in the second week, 70% in the third week, 60% in the fourth week, and then kept at 60% of the normal level for the remaining 56 weeks? Or another 60 weeks; (3) Calorie Restriction-Recovery group (CR-R group) involved CR for 52 weeks and then access to food *ad libitum*. A 24-h urine sample was collected from each rat at 0, 1, 2, 3, 6, 10, 27, 42, 51, 54 and 60 weeks of CR. Contents of colon and ileum, feces, liver, and blood were collected when rats were sacrificed, stored at −80 °C storage for future metabolomics analysis. Further, when the rats were sacrificed after 60 weeks of CR, ileum contents were collected for further 16s rRNA sequencing. Cecal and colon contents were also collected for further *Lactobacillus* culture study. For Metagenomic sequencing, 5 ileum content samples in each group were randomly selected and mixed into a pooled sample. For metatranscriptome sequencing, 5 samples of cecal contents in each group were randomly chosen.

For the short-term calorie restriction, 40d-CR study, after 4 weeks of acclimatization in metabolic cages, 20 eight-week-old male Wistar rats were randomly divided into C group and CR group. The CR treatment only lasted for 40 days. A 24-h urine sample was collected from each rat at 0 and 40 days of CR. Contents of colon and ileum, serum, feces and liver were collected when the rats were sacrificed after 40 days of CR.

For the CR-antibiotic treatment experiment, after 4 weeks of acclimatization in metabolic cages, rats were randomly divided into C group, CR group (CR treatment for 26 weeks); and (3) Vancomycin-CR group (CRV group), which were treated with Vancomycin solution (200mg/kg.d, *ip*) for 5 days after CR for 26 weeks. Urine and gut contents were collected after the rats were sacrificed at the end of experiment.

For CR-Mice study, after 4 weeks of acclimatization in metabolic cages, 20 eight-week-old C57BL/6 mice were randomly divided into C group, CR group (CR treatment for 26 weeks).

Urine and serum samples were centrifuged at 1409 *g* for 10 min at room temperature, and the resulting supernatants were used for experiments. Urine, serum, liver, and intestinal contents and feces were stored at −80 °C until assay.

### GC/MS Sample Preparation, Derivatization, and Spectral Acquisition

Urine samples were prepared according to published methods with minor modifications^48^. Briefly, Pooled quality control (QC) samples were prepared by mixing 20 μL of each single sample. 100µL of urine sample and 10µL of ureases (30U/10µL) were vortexed for 10s. The mixed solution was incubated at 37°C for 15min before mixing with the standard solutions (10 μL L-2-chlorophenylalanine in water, 0.1 mg/mL; 10μL heptadecanoic acid in methanol, 1 mg/mL) and 300µL methanol. After being vortexed for 30s, samples were centrifuged at 13,200 rpm for 5 min at −4°C and an aliquot of the resulting 300μL of supernatant was dried in a vacuum dryer at room temperature. The residue was derivatized using a two-step procedure: 80μL methoxyamine (15 mg/mL in pyridine) was added to the vial and incubated at 30 °C for 90 min, followed by an incubation with 80μL of N,O-Bis(trimethylsilyl)trifluoroacetamide (BSTFA) (1% TMCS) at 70 °C for 60 min.

Serum samples were prepared according to published methods with minor modifications. ^49^ Pooled quality control (QC) samples were prepared by mixing 20 μL of each serum sample. A 50 μL aliquot of serum sample was spiked with two internal standards (10 μL of L-2-chlorophenylalanine in water, 0.3 mg/mL; 10 μL of heptadecanoic acid in methanol, 1 mg/mL) and vortexed for 10s. The mixed solution was extracted with 175 μL of methanol/chloroform (3:1) and vortexed for 30 s. AGer the samples were stored for 10 min at −20 °C, they were centrifuged at 8000 rpm for 10 min. An aliquot of the 200 μL supernatant was transferred to a glass sampling vial and was vacuum-dried at room temperature. The residue was derivatized using a two-step procedure: 80μL methoxyamine (15 mg/mL in pyridine) was added to the vial and incubated at 30 °C for 90 min, followed by an incubation with 80μL BSTFA (1% TMCS) at 70 °C for 60 min. When the reaction was finished, the samples were placed at room temperature for 1 h waiting for GC-TOFMS analysis.

Gut contents and gut bacteria culture medium samples were prepared according to published methods with minor modifications.^50^ Gut contents water samples were mixed using 30mg of thawed samples and 100μL water followed by short-time vortexing at room temperature. A 3-fold volume of ice-cold methanol chloroform solvent (3:1) was added to the gut contents samples and gut bacteria culture medium samples. The mixture was vortexed strongly and subsequently centrifuged at 14 000 rpm at 4 °C for 15 min. Pooled quality control (QC) samples were prepared by mixing 20 μL of each supernatant of the previous mixture. 300 μL of supernatant were freeze-dried, and subsequently, 80 μL of methoxylamine solution (15 mg/mL in pyridine) was added to each vial. The resultant mixture was vigorously vortex-mixed for 1 min and reacted at 37 °C for 24 h. Eighty microliters of BSTFA (with 1% TMCS) were added into the mixture which was then derivatized at 70 °C for 90 min, and vortexed just prior to injection onto the GC/TOFMS.

Each 1μL aliquot of the derivatized solution was injected in splitless mode into an Agilent 6890N gas-chromatograph coupled with a Pegasus HT time-of-flight mass spectrometer (GC/TOF, Leco Corp., St. Joseph, MI). Separation was achieved on a DB-5ms capillary column (30m × 250μm i.d., 0.25μm film, (5%-phenyl)-methylpolysiloxane bonded and cross-linked; Agilent J&W Scientific, Folsom, CA) with helium as the carrier gas at a constant flow rate of 1.0 mL/ min. The temperature of injection, transfer interface, and ion source was set to 270, 270, and 220 °C, respectively. The GC temperature programming was set to 2 min isothermal heating at 80 °C, followed by 10 °C/min oven temperature ramp to 140 °C, 4 °C/min to 240 °C, and 25 °C/min to 290 °C, and a final 4.5 min maintained temperature at 290 °C. Electron impact ionization (70 eV) at full scan mode (m/z 30−600) was used with an acquisition rate of 20 spectra/s in the TOFMS setting.

### GC/MS Data Reduction and Pattern Recognition

The peak information from GC/MS data was converted, normalized and prepared prior to the multivariate analysis. The PCA model was used to identify the urinary metabolites differentially produced by caloric restriction and the refeeding procedure.

Fold changes of the arithmetic mean values (each group/control group) and p-values were calculated by using the Kruskal-Wallis test of all the differentially expressed metabolites along with standard compounds. In addition to this nonparametric test, classical one-way analysis of variance (ANOVA) was also used to judge the statistical significance of the results. The critical p-value of both tests was set to 0.05 for this study.

### Ileum contents microbial taxonomy by 16S rRNA sequencing

Genome DNAs were extracted from ileum samples using the QIAamp DNA stool mini kit (QIAGEN, cat#51504) following the manufacturer’s recommendations and the DNA samples were stored at −20 °C until sequencing. Next generation sequencing library preparations and IlluminaMiSeq sequencing were conducted at GENEWIZ, Inc. (Beijing, China). DNA samples were quantified using a Qubit 2.0 Fluorometer (Invitrogen, Carlsbad, CA) and DNA quality was checked on a 0.8% agarose gel. DNA samples (5-50 ng) were used to generate amplicons using a MetaVx™ Library Preparation kit (GENEWIZ, Inc., South Plainfield, NJ, USA). A panel of proprietary primers was designed to anneal to the relatively conserved regions bordering the V3, V4, and V5 hypervariable regions. The v3 and v4 regions were amplified using forward primers containing the sequence “CCTACGGRRBGCASCAGKVRVGAAT” and reverse primers containing the sequence “GGACTACNVGGGTWTCTAATCC”. The v4 and v5 regions were amplified using forward primers containing the sequence” GTGYCAGCMGCCGCGGTAA” and reverse primers containing the sequence “CTTGTGCGGKCCCCCGYCAATTC”. Besides the 16S target-specific sequence, the primers also contain adaptor sequences allowing uniform amplification of the library with high complexity ready for downstream NGS sequencing on IlluminaMiseq.

DNA libraries were validated using an Agilent 2100 Bioanalyzer (Agilent Technologies, Palo Alto, CA, USA), and quantified by Qubit and real time PCR (Applied Biosystems, Carlsbad, CA, USA). DNA libraries were multiplexed and loaded on an IlluminaMiSeq instrument according to manufacturer’s instructions (Illumina, San Diego, CA, USA). Sequencing was performed using a 2×250 or 2×300 paired-end (PE) configuration; image analysis and base calling were conducted by the MiSeq Control Software (MCS) on the MiSeq instrument.

The QIIME data analysis package was used for 16S rRNA data analysis. The forward and reverse reads were joined and assigned to samples based on barcode and truncated by cutting off the barcode and primer sequence. Quality filtering on joined sequences was performed and sequence which did not fulfill the following criteria were discarded: sequence length <200bp, no ambiguous bases, mean quality score ≥ 20. Then the sequences were compared with the reference database (RDP Gold database) using UCHIME algorithm to detect chimeric sequences which were removed.

The effective sequences were used in the final analysis. Sequences were grouped into operational taxonomic units (OTUs) using the clustering program VSEARCH(1.9.6) against the Silva 119 database pre-clustered at 97% sequence identity. The Ribosomal Database Program (RDP) classifier was used to assign taxonomic category to all OTUs at confidence threshold of 0.8. The RDP classifier uses the 16SrRNA RDP database which has taxonomic categories predicted to the species level.

Sequences were rarefied prior to calculation of alpha and beta diversity statistics. Alpha diversity indexes were calculated in QIIME from rarefied samples using for diversity the Shannon index, for richness the Chao1 index. Beta diversity was calculated using weighted and unweighted UniFrac and principal coordinate analysis (PCoA) performed. Unweighted Pair Group Method with Arithmetic mean (UPGMA) tree from beta diversity distance matrix was built.

The sequences were aligned using FLASH (V1.0.3)^51^ and the unaligned ones were discarded. The delineation of OTUs was conducted using QIIME at 97% cutoff^52^. The length of the sequence fragments used for the analysis was from 178 to 266 nucleotides (without primer and barcode). The alpha and beta diversities were performed using mothur^53^.

We performed LEfSe analysis (http://huttenhower.sph.harvard.edu/galaxy) ^54^ which can discover high-dimensional biomarker and identify genomic features (genes, pathways or taxa) characterizing the differences between two or more biological conditions. The differential features were identified on the OTU level. The treatment groups were used as the class of subjects (no subclass). LEfSe analysis was performed under the following conditions: (1) the alpha value for the factorial Kruskal-Wallis test among classes is <0.01 and (2) the threshold on the logarithmic LDA score for discriminative features is >2.0. The data of relative abundance in each sample for each OTU selected by LEfSe was shown in a heatmap by using power transformations.

### Taxonomic and functional profiling of metagenomic samples

#### Library Preparation and Sequencing

Genome DNAs were extracted from pooled rat ileum contents samples using a QIAamp DNA stool mini kit (QIAGEN, cat#51504) following the manufacturer’s recommendations and the DNA samples were stored at −20 °C until sequencing. Next generation sequencing library preparations were constructed following the manufacturer’s protocol (NEBNext^®^ Ultra™ DNA Library Prep Kit for Illumina^®^). For each sample, 2 µg of genomic DNA was randomly fragmented to <500 bp by sonication (Covaris S220). The fragments were treated with End Prep Enzyme Mix to repair both ends and to add a dA-tailing in one reaction, followed by a T-A ligation to add adaptors to both ends. Size selection of Adaptorligated DNA was then performed using AxyPrep Mag PCR Clean-up (Axygen) and fragments of ~410 bp (with the approximate insert size of 350 bp) were recovered. Each sample was then amplified by PCR for 8 cycles using P5 and P7 primers, with both primers carrying sequences which can anneal with flow cell to perform bridge PCR and P7 primer carrying a six-base index allowing for multiplexing. The PCR products were cleaned up using AxyPrep Mag PCR Cleanup (Axygen), validated using an Agilent 2100 Bioanalyzer (Agilent Technologies, Palo Alto, CA, USA), and quantified by using a Qubit2.0 Fluorometer (Invitrogen, Carlsbad, CA, USA). The libraries with different indexes were multiplexed and loaded on an IlluminaHiSeq instrument according to manufacturer’s instructions (Illumina, San Diego, CA, USA). Sequencing was carried out using a 2×150 paired-end (PE) configuration; image analysis and base calling were conducted by the HiSeq Control Software (HCS) + OLB + GAPipeline-1.6 (Illumina) on the HiSeq instrument.

#### Data Analysis

Raw shotgun sequencing reads were trimmed using cutadapt (v1.9.1). Low-quality reads, Nrich reads and adapter-polluted reads were removed. Then host contamination reads were removed. Samples were each assembled *de novo* to obtain separate assemblies. Whole genome *de novo* assemblies were performed using SOAPdenovo(v2)with different k-mer. The best assembly result of Scaffold, which has the largest N50, was selected for the subsequent analysis. CD-HIT was used to cluster scaftigs derived from assembly with a default identity of 0.95.

In order to analyze the relative abundance of scaftigs in each sample, paired-end clean reads were mapped to assembled scaftigs using the Burrows-Wheeler Aligner (BWA version 0.7.12) to generate read coverage information for assembled scaftigs. Paired forward and reverse read alignments were generated in the SAM format using the BWA-SAMPE algorithm with default parameters. The mapped reads counts were extracted using SAMtools 0.1.17. The corresponding scaftigs were mapped to the mass of Bacteria, Fungi, Archaea and Viruses data extracted from the NT database of NCBI. LCA algorithm (Lowest common ancestor, applied in MEGAN software system) was used to ensure the annotation significance by picking out the lowest common classified ancestor for final display.

Genes were predicted using MetaGeneMark, and BLASTP was used to search the protein sequences of the predicted genes with the NR database, CAZy database, eggNOG database and KEGG database with E < 1e-5. To determine the similarity or difference of taxonomic and functional components between different samples, relative Clustering analysis and Principal Component Analysis (PCA) were performed. Meanwhile, there were a series of advanced analysis items available to explore the environmental samples, such as LEfSe, significant difference, CCA/RDA, NMDS, prediction of secreted protein an annotation of CARD, VFDB, MvirDB, PHI, TCDB.

### Taxonomic and functional profiling of Metatranscriptome

Extraction of total RNA was performed using TRIzol Reagent (Cat. No. 10296010; Invitrogen, USA). The rRNA sequence specific probe was hybridized with the total RNA, then the rRNA / probe complex was removed by magnetic beads and further purified by ethanol precipitation. Complementary DNA (cDNA) was obtained using TruSeq Stranded mRNA LT Sample Prep Kit (Illumina), which includes RNA fragmentation, cDNA synthesis and cDNA library construction. An Agilent Bioanalyzer was used to do a 2100 quality inspection on the library. The qualified library should have a single peak and no joint. Then the library was quantified using a Promega quantifluor. The concentration of the qualified library should be more than 2nM after calculation. The qualified library was sequenced 2 × 150 bp. The library (index non repe) that needs to be on the computer was diluted by gradient. Then the samples were mixed according to the required data volume proportion. The mixed library was transformed into a single chain and sequenced using Illumina HiSeq X-ten platform (Illumina, USA) with PE150 strategy at Personal Biotechnology Co., Ltd. (Shanghai, China). The nucleotide sequences determined in this study have been deposited under accession numbers.

Raw sequencing reads were processed to obtain quality-filtered reads for further analysis. First, sequencing adapters were removed from sequencing reads using Cutadapt (v1.2.1) (Martin 2011). Second, low quality reads were trimmed by using a sliding-window algorithm. Third, reads were aligned to the host genome using BWA (http://bio-bwa.sourceforge.net/) (Li and Durbin 2009) to remove host contamination. Once quality-filtered reads were obtained, they were *de novo* assembled to construct the metagenome for each sample by IDBA-UD (Iterative De Bruijn graph Assembler for sequencing data with highly Uneven Depth) (Peng, Leung et al. 2012). All coding regions (CDS) of metagenomic scaffolds longer than 300 bp were predicted by MetaGeneMark (http://exon.gatech.edu/GeneMark/metagenomew) (Zhu, Lomsadze et al. 2010).

CDS sequences of all samples were clustered by CD-HIT (Fu, Niu et al. 2012) at 90% protein sequence identity to obtain a non-redundant gene catalog. Gene abundances in each sample was estimated by soap.coverage (http://soap.genomics.org.cn/) based on the number of aligned reads. The lowest common ancestor taxonomy of the non-redundant genes was obtained by aligning them against the NCBI-NT database by BLASTN (e value < 0.001). Similarly, the functional profiles of the non-redundant genes were obtained by annotating against the GO, KEGG, EggNOG and CAZy databases, respectively, by using the DIAMOND (Buchfink, Xie et al. 2015) alignment algorithm. Based on the taxonomic and functional profiles of non-redundant genes, LEfSe (Linear discriminant analysis effect size) was performed to detect differentially abundant taxa and functions across groups using the default parameters (Segata, Izard et al. 2011). Beta diversity analysis was performed to investigate the compositional and functional variation of microbial communities across samples using Bray-Curtis distance metrics (Bray and Curtis 1957) and visualized via principal coordinate analysis (PCoA), nonmetric multidimensional scaling (NMDS) and unweighted pair-group method with arithmetic means (UPGMA) hierarchical clustering (Ramette 2007).

Through the galaxy online analysis platform (http://huttenhower.sph.harvard.edu/galaxy/), we submitted the abundance spectrum of the underlying functional groups annotated by the macro transcriptome samples in each functional database, or the composition of each level for LEfSe analysis.

The functional annotation results of each Unigene in the KEGG database, combined with its corresponding expression spectrum, can be used to analyze the expression distribution of samples in the KO level.

### qPCR for Glyoxylate cycle gene-quantitation

Six ileum content samples were randomly collected from each group(C and CR) of 60w-CR. DNA was extracted using a QIAamp^®^ DNA Stool Mini Kit (Qiangen, Germany) from samples of each rat (C and CR). Primers were selected from the NGS of environmental microbes’ RNA sequences (Supplementary Table S3). Primers D1 (icl-F 5’-TGACCAGTGTCGGCATCGGC-3’) and D2 (icl-R 5’-CGCATGAGCCCTGAGGAGCG-3’) and D3 (ms-F 5’-GCCTGGCTCGTTGGAGGTCG-3’) and D4 (ms-R 5’-CGTGAGCACCAGGCCAAGGG-3’). DNA was detected spectrophotometrically at 280 nm using a Nano Drop 2000C (Thermo, USA). qPCR reaction reagent using Quanti Nova SYBR^®^ Geen PCR kit(Qiangen, Germany) and qPCR reactions containing 1ul template,6ul ddH_2_0, 1ul ROX, 10ul 2×SYBR real-time PCRmix and 1ul primer forward and reverse(10uM), respectively in a 20ul total volume. Thermal cycling(ABI Step one plus 7900, USA) were using 95°C for 10min, 40× (95°C for 30 sec, 60°C for 30 sec).

### RT-PCR for hepatic TCA cycle gene-quantitation

#### RNA extraction and cDNA synthesis

Eight liver tissue samples were randomly collected from each group(C and CR) of 60w-CR. RNA was extracted using Trizol (Invitrogen) and was reverse transcribed into cDNA using a Super RT Kit (TaKaRa, China) at 42°C using the protocols provided by the manufacturer. The primers are listed (Supplementary Table S4 GAPDH as the housekeeping gene).cDNA was diluted 1:10 using ddH_2_0.

#### Quantitative PCR

Each RT-PCR reaction was run on an ABI Step one plus 7900(USA).Every 20ul volume system reaction consisted of 10ul 2×SYBR real-time PCR mix, 0.4ul primer forward and reverse (10uM), 1ul cDNA, along with addition of ddH20 to 20ul. The thermal cycling were 95°C for 5min and 40 cycles of amplification (95°C 15sec, 60°C 30sec). Each reaction was run with a melting curve to identify each pair of primers with only one significant peak.

### Acetate intervention on gut microbiota culture medium

In 60w-CR study, five rats from each group were randomly selected to collect 200mg of fresh ileum contents. Then M9 medium 3800μL, 1000 × cgcl_2_ 3.8 μL, 500 × MgSO_4_ 7.6 μL and glucose (2%) 76 μL were added into the samples and mixed. Glacial acetic acid (5.0 μL) was added to the culture medium to make a final concentration of 25 mM. Cultures were incubated anaerobically at 37°C. At 0, 1, 3 and 6 hours, an aliquot of 100 μL of conditioned media for each sample was taken out for bacteria isolation and identification, and an aliquot of 200 μL for each sample was taken out for metabolomics analysis. The aforementioned 100 μL culture medium was diluted with M9 solvent to 10^−5^, and scribed onto the M9 medium plate containing sodium acetate (25mM), and cultured at 37 °C anaerobically for 2 days. The PCR products purified from each strain were taken and sequenced with abi3730-xl sequencer. The sequence obtained was searched and compared in NCBI https://blast.ncbi.nlm.nih.gov/Blast.cgi?PROGRAMblastn&PAGE_TYPEBlastSearch&LINK_LOCblasthome and the sequence with the largest similarity was selected as the result of species identification.

### Lactobacillus proliferation experiment in the gut microbial starvation culture of different intestinal segments

#### *Lactobacillus* murinus isolation and culture

In 60w-CR study, 3 rats from the control group were randomly selected to collect fresh gut contents. Ileum, cecum and colon contents were put into 2mL EP tube respectively, sterile Luria BERTANI (LB) medium was added, and samples were mixed well and then filtered with 40 μM filter screen to remove the large particles in the bacterial solution. The filtrate volume was collected in a 50ml EP tube. Then, the filtrate was filtered through a 0.22 μM filter membrane, and the intestinal bacteria on the filter membrane were collected again in a new 50mL EP tube. The metabolites of the original gut contents were filtered out to obtain pure intestinal flora, and new LB medium was added to 48mL. The medium was divided into 1.5mL per EP tube, then transferred to an anaerobic tank (N_2_: H_2_: CO_2_ = 85:10:5) to replace the gas, and cultured at 37 °C for 0h, 3h, 6h, 8h, 24h, 48h and 72h respectively to obtain the metabolites of gut bacteria at different times. The bacterial medium at each time point were taken out in turn, and were filtered with 0.22 μ M filter membrane in in sterile environment, to obtain pure gut bacterial metabolite solution. Then 900 μL of the filtrate was added to 80 μL *Lactobacillus murinus* that was isolated and cultured previously, and then placed in an anaerobic tank for 24 hours. Then the culture medium was filtered to remove *Lactobacillus*. The filtrate 500 μL were taken and added with 500 μL pre-cooled methanol, mix well and put it into −80 °C refrigerator immediately for preservation. Preservation solution (100 μL) was taken out for measurement of its OD value using an ultraviolet spectrophotometer.

### Western blot analysis

Cells proteins and tissue were lysed with RIPA buffer containing protease and phosphatase inhibitors, and analyzed by SDS–PAGE and western blot. Proteins were detected using specific primary antibodies directed against LC3A/B (#12741, 1:1000, Cell Signaling Technology, USA), Phospho-AMPKα(#2535, 1:1000, Cell Signaling Technology, USA), AMPKα(#2603,1:1000, Cell Signaling Technology, USA), SQSTM1/p62 (#5114,1:1000, Cell Signaling Technology, USA), β-TransGen Biotech, China) for 1 h. The Sigma actin(, 1:3000, Cell Signal Technology, USA),Atg7 (8558, 1:1000, Cell Signal Technology, USA), The protein expression levels were normalized with β-actin (1:3000, TransGen Biotech, China). After washing with TBST, the membrane was incubated with HRP-conjugated secondary antibody (1:10000, l was detected by using Odyssey9120 system.

### Primary hepatocyte isolation, cell culture and treatment

Methods Primary hepatocyte isolation, culture and treatment Primary rat hepatocytes were isolated following a two-step *in situ* collagenase perfusion method as described previously^55^. Hepatocytes were seeded at subconfluence (0.5 × 106/ ml) in Williams E Medium (Gibco, Life Technologies, Carlsbad, CA) supplemented with 10 % (v/v) fetal bovine serum (Hyclone, Logan, Utah), 0.1 μM dexamethasone, 10 μg/ml insulin, 2 mM L-glutamine, and 100 μg/ml penicillin and streptomycin in collagen I-coated cell plate. Four hour after incubation, the cells were washed and incubated in serum-free Williams E Medium. Hepatocytes were cultured at 37 °C, 5 % CO2, and 95% relative humidity.

Malate solution was adjusted to pH 6-7 with sodium hydroxide to the series concentrations. For western blot analysis, hepatocytes, HCT116, NCM460 cell lines and primary rat hepatocytes were seeded in 6-well plates and treated with malate solution at using serial dilution for 12 hours.

## ACKNOWLEDGMENTS

This work was financially supported by the National Basic Research Program of China (2012CB910102), Shanghai Jiao Tong University Biomedical Engineering Cross Research Foundation (YG2015MS15, YG2016MS40) and National Nature Science Foundation of China (30901997).

