## Supplementary material for "Caloric Restriction Remodels Energy Metabolic Pathways of Gut Microbiota and Promotes Host Autophagy": SI-Figures and TableS1-S2

**Table S1** The diet composition of standard chow

| Diet composition (%) | |
|---|---|
| Moisture | ≤ 10 |
| Crude protein | ≥ 18 |
| Crude fat | ≥ 4 |
| Crude fiber | ≤ 5 |
| Crude ash | ≤ 8 |
| Calcium | 1.0 – 1.8 |
| Total phosphorus | 0.6 – 1.2 |
| Lysine | ≥ 0.82 |
| Methionine + Cystine | ≥ 0.53 |
| Sodium chloride | 0.4 |

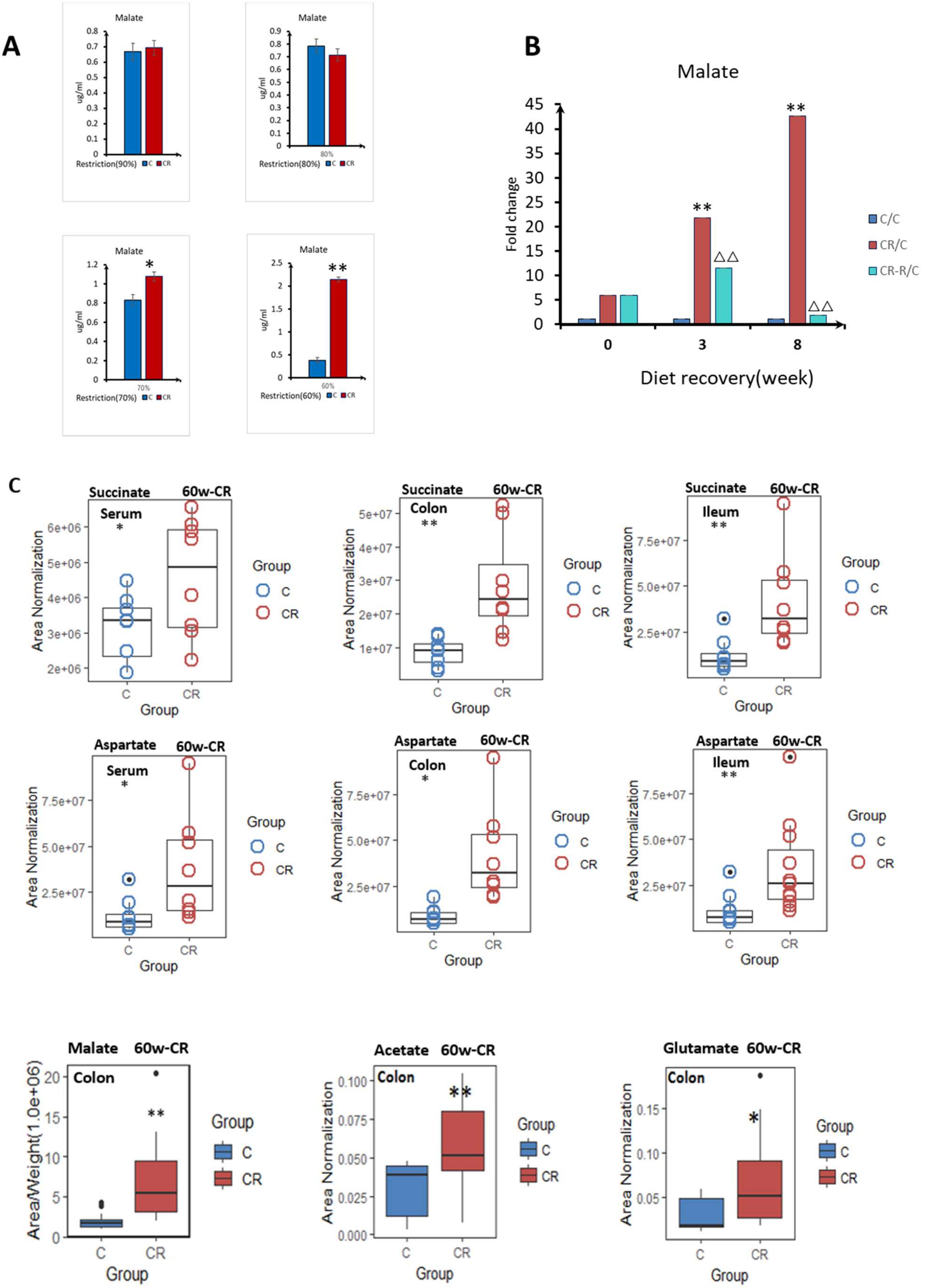
A
Malate
ug/ml
Restriction(90%)
C
CR
Malate
ug/ml
80%
Restriction(80%)
C
CR
Malate
*
ug/ml
70%
Restriction(70%)
C
CR
Malate
**
ug/ml
60%
Restriction(60%)
C
CR
B
Malate
Fold change
**
**
△△
△△
C/C
CR/C
CR-R/C
0
3
8
Diet recovery(week)
C
Succinate 60w-CR
Serum
*
Area Normalization
Group
C
CR
Succinate 60w-CR
Colon
**
Area Normalization
Group
C
CR
Succinate 60w-CR
Ileum
**
Area Normalization
Group
C
CR
Aspartate 60w-CR
Serum
*
Area Normalization
Group
C
CR
Aspartate 60w-CR
Colon
*
Area Normalization
Group
C
CR
Aspartate 60w-CR
Ileum
**
Area Normalization
Group
C
CR
Malate 60w-CR
Colon
**
AreaWeight(1.0e+06)
Group
C
CR
Acetate 60w-CR
Colon
**
Area Normalization
Group
C
CR
Glutamate 60w-CR
Colon
*
Area Normalization
Group
C
CR

Fig.S1 Variations of malate levels during 60w-CR.

A) Malate changes in varying degrees of diet restriction from 90% to 60%; B) urine malate level 3 weeks and 8 weeks after diet recovery (compared with C group); C) Succinate and aspartate variances in serum, colon and ileum after 60w-CR; *, T test($p<0.05$),**,T test($p<0.01$) (compared with CR group:△, T test($p<0.05$),△△,T test($p<0.01$)

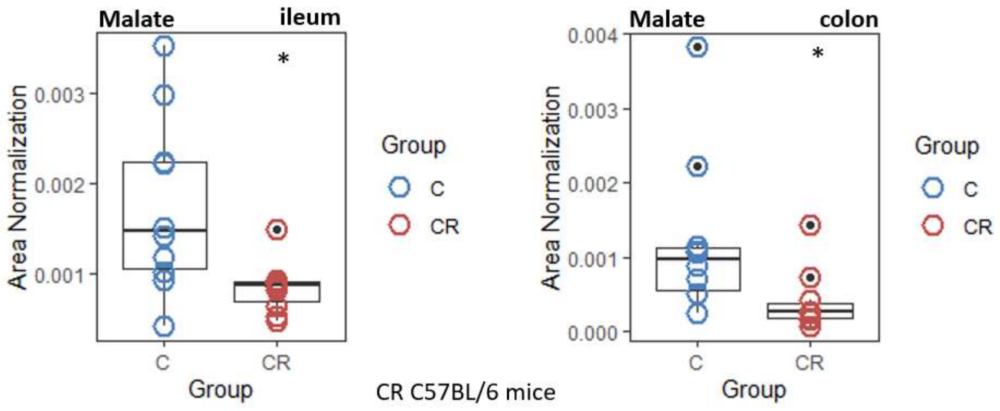


Fig.S2 Malate changes in gut contents from CR C57BL/6 mice for 26 weeks. *, T test (p<0.05)

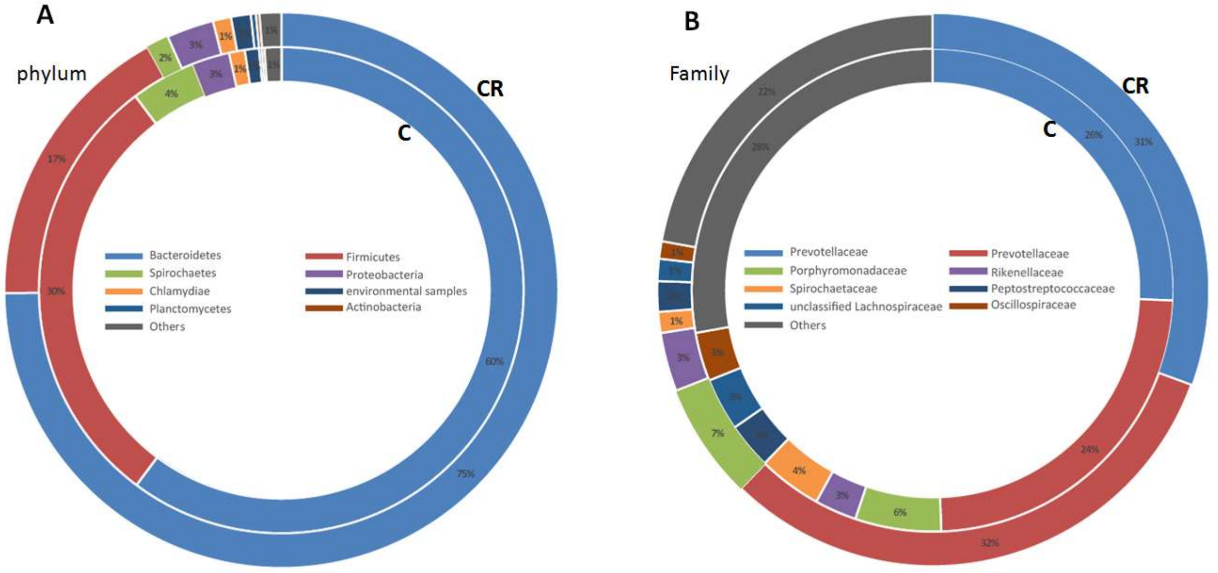


Fig.S3 The characteristic changes of ileum flora induced by 60w-CR.

A) 16s Microbiome structure comparison between C and CR groups at phylum (B) and genus level

A

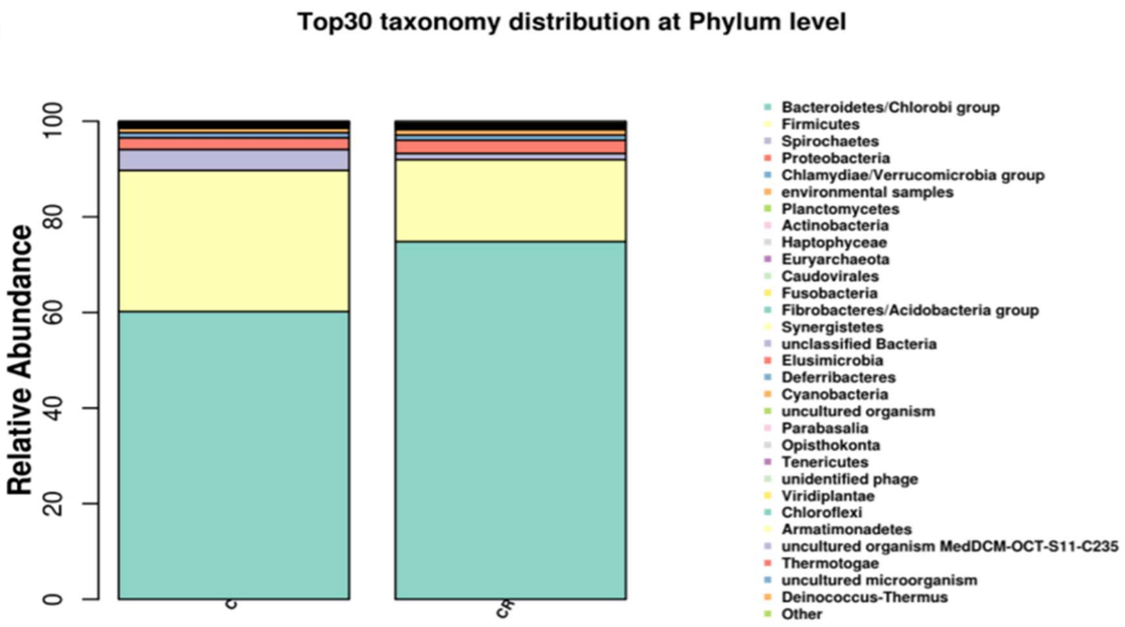
Top30 taxonomy distribution at Phylum level
Relative Abundance
0
20
40
60
80
100
C
CR
Bacteroidetes/Chlorobi group
Firmicutes
Spirochaetes
Proteobacteria
Chlamydiae/Verrucomicrobia group
environmental samples
Planctomycetes
Actinobacteria
Haptophyceae
Euryarchaeota
Caudovirales
Fusobacteria
Fibrobacteres/Acidobacteria group
Synergistetes
unclassified Bacteria
Elusimicrobia
Deferribacteres
Cyanobacteria
uncultured organism
Parabasalia
Opisthokonta
Tenericutes
unidentified phage
Viridiplantae
Chloroflexi
Armatimonadetes
uncultured organism MedDCM-OCT-S11-C235
Thermotogae
uncultured microorganism
Deinococcus-Thermus
Other

B

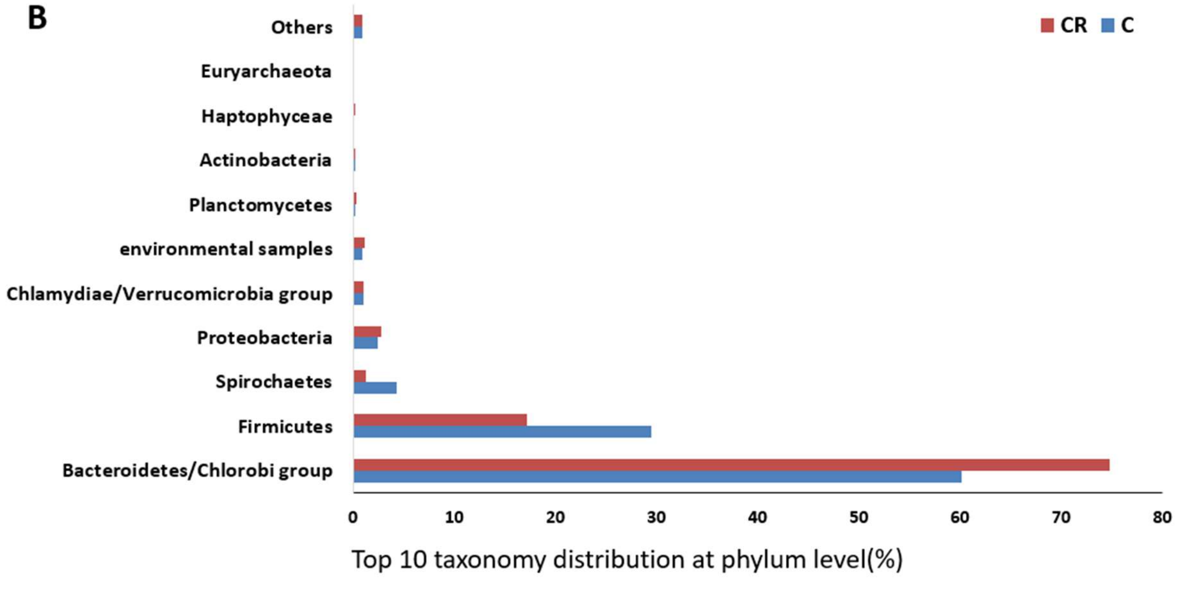
CR
C
Others
Euryarchaeota
Haptophyceae
Actinobacteria
Planctomycetes
environmental samples
Chlamydiae/Verrucomicrobia group
Proteobacteria
Spirochaetes
Firmicutes
Bacteroidetes/Chlorobi group
0
10
20
30
40
50
60
70
80
Top 10 taxonomy distribution at phylum level(%)

C

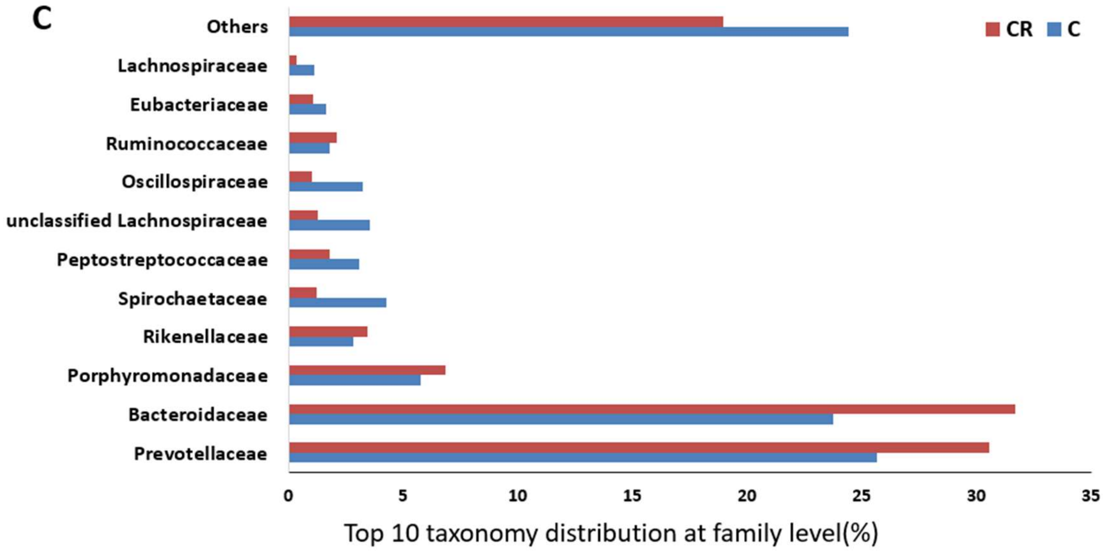


D

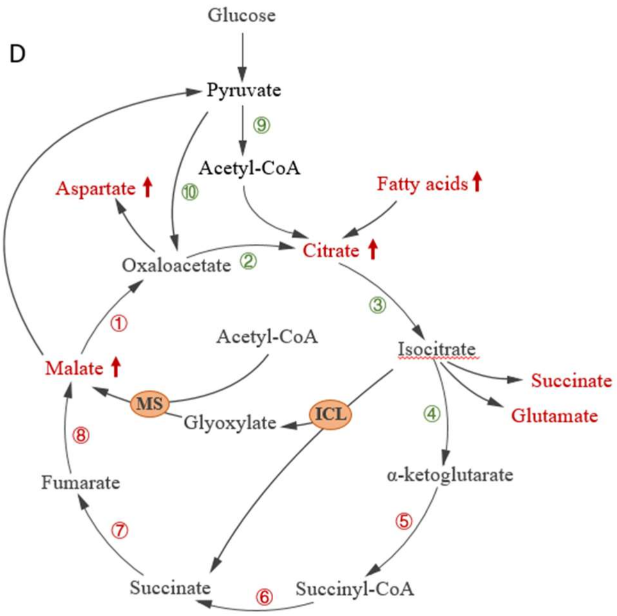


| Number | Enzyme name | Entry | Fold change |
|---|---|---|---|
| 1 | Malate dehydrogenase | EC:1.1.1.37 、 EC:1.1.5.4 | 1.36 |
| 2 | Citrate synthase | EC:2.3.3.1 | 0.67 |
| 3 | Aconitate hydratase | EC:4.2.1.3 | 0.76 |
| 4 | Isocitrate dehydrogenase | EC:1.1.1.41 、 EC:1.1.1.42 | 0.65 |
| 5 | 2-oxoglutarate dehydrogenase | EC:1.2.4.2 、 EC:2.3.1.61 | 1.35 |
| 6 | Succinyl-CoA synthetase | EC:6.2.1.4 、 EC:6.2.1.5 | 1.28 |
| 7 | Succinate dehydrogenase / fumarate reductase | EC:1.3.5.1 、 EC:1.3.5.4 | 1.38 |
| 8 | Fumarate hydratase | EC:4.2.1.2 | 1.09 |
| 9 | Pyruvate dehydrogenas | EC:1.2.4.1 、 EC:2.3.1.12 | 0.38 |
| 10 | Pyruvate carboxylase | EC:6.4.1.1 | 1.04 |

E

| Gene_ID | Genus | Gene_ Name | Definition | EC | Pathway |
|---|---|---|---|---|---|
| **gene_168766** | Prevotellaceae | gph | phosphoglycolate phosphatase | EC:3.1.3.18 | ko00630:Glyoxylate and dicarboxylate metabolism; |
| **gene_107514** | Bacteroidaceae | oxc | oxalyl-CoA decarboxylase | EC:4.1.1.8 | ko00630:Glyoxylate and dicarboxylate metabolism; |
| **gene_191922** | Bacteroidaceae | gph | phosphoglycolate phosphatase | EC:3.1.3.18 | ko00630:Glyoxylate and dicarboxylate metabolism; |
| **gene_260363** | Rikenellaceae | gph | phosphoglycolate phosphatase | EC:3.1.3.18 | ko00630:Glyoxylate and |

| | | | | | |
|---|---|---|---|---|---|
| | | | | | dicarboxylate metabolism; |
| **gene_243309** | Prevotellaceae | gph | phosphoglycolate phosphatase | EC:3.1.3.18 | ko00630:Glyoxylate and dicarboxylate metabolism; |
| **gene_200884** | Prevotellaceae | gph | phosphoglycolate phosphatase | EC:3.1.3.18 | ko00630:Glyoxylate and dicarboxylate metabolism; |
| **gene_304722** | Bacteroidaceae | oxc | oxalyl-CoA decarboxylase | EC:4.1.1.8 | ko00630:Glyoxylate and dicarboxylate metabolism; |
| **gene_378590** | Prevotellaceae | gph | phosphoglycolate phosphatase | EC:3.1.3.18 | ko00630:Glyoxylate and dicarboxylate metabolism; |
| **gene_56580** | Bacteroidaceae | gph | phosphoglycolate phosphatase | EC:3.1.3.18 | ko00630:Glyoxylate and dicarboxylate metabolism; |
| **gene_206865** | Parasutterella | glcD | glycolate oxidase | EC:1.1.3.15 | ko00630:Glyoxylate and dicarboxylate metabolism; |
| **gene_414359** | Parasutterella | gph | phosphoglycolate phosphatase | EC:3.1.3.18 | ko00630:Glyoxylate and dicarboxylate metabolism; |
| **gene_248160** | Parasutterella | fdoG, fdfH | formate dehydrogenase major subunit | EC:1.2.1.2 | ko00630:Glyoxylate and dicarboxylate metabolism |

Fig.S4 Results of metagenomic sequencing
A)Taxonomy assignment of phylum Level, B) Top 10 taxonomy distribution at phylum and C)genus level, in metagenomic results; D) TCA cycle and glyoxylate cycle related differential expressed genes; E) A List of top 10 glyoxylate cycle pathways associated genes and bacteria genus in metagenomic results.
CR-dependent variation of malate and glyoxylate cycle related bacteria.

A

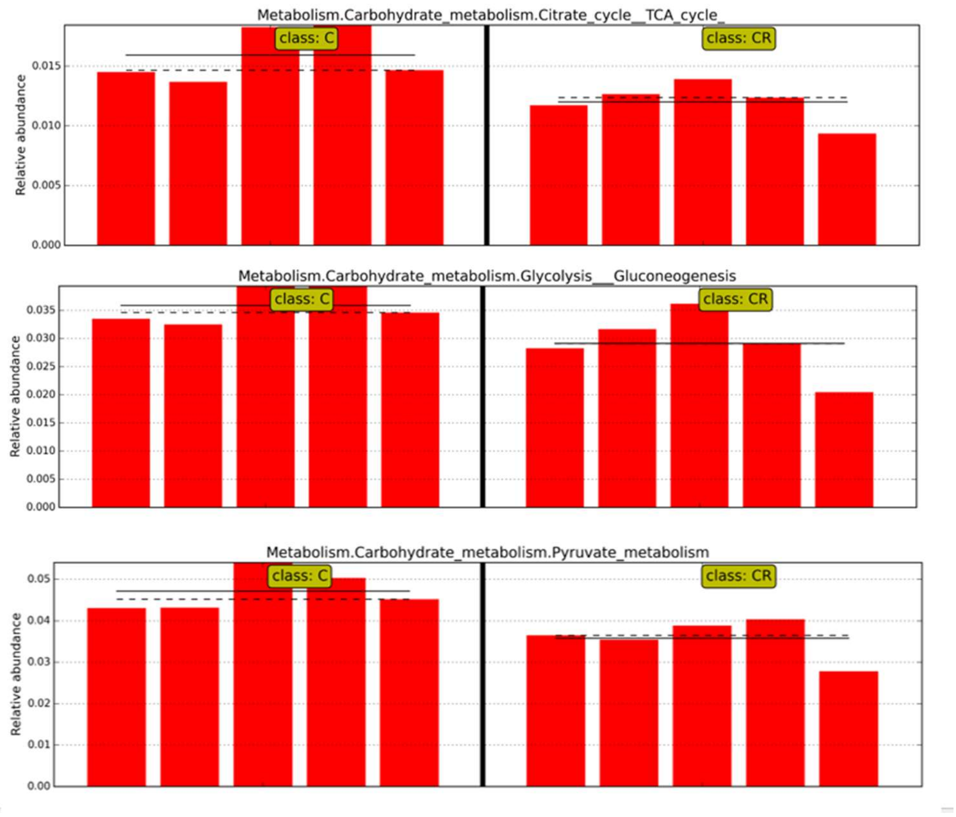
Metabolism.Carbohydrate_metabolism.Citrate_cycle__TCA_cycle_
class: C
class: CR
Relative abundance
0.000
0.005
0.010
0.015
Metabolism.Carbohydrate_metabolism.Glycolysis__Gluconeogenesis
class: C
class: CR
Relative abundance
0.000
0.005
0.010
0.015
0.020
0.025
0.030
0.035
Metabolism.Carbohydrate_metabolism.Pyruvate_metabolism
class: C
class: CR
Relative abundance
0.00
0.01
0.02
0.03
0.04
0.05

B

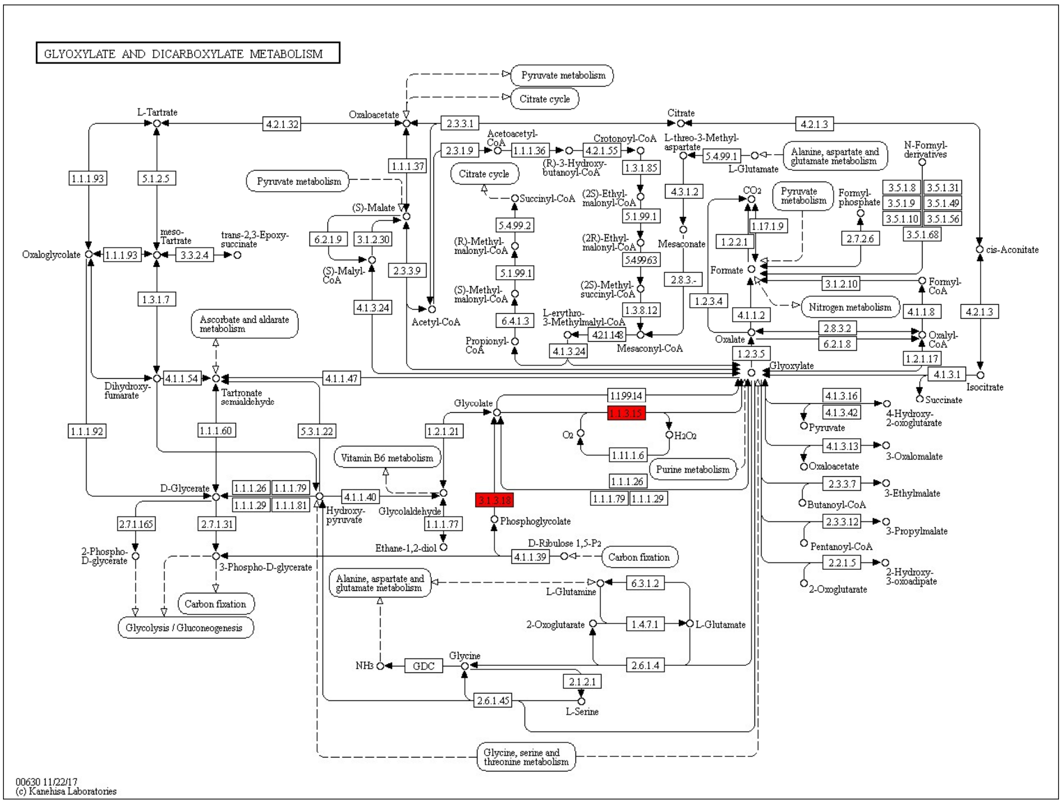
GLYOXYLATE AND DICARBOXYLATE METABOLISM
Pyruvate metabolism
Citrate cycle
L-Tartrate
4.2.1.32
Oxaloacetate
2.3.3.1
Citrate
4.2.1.3
Acetoacetyl-CoA
Crotonoyl-CoA
L-threo-3-Methyl-aspartate
N-Formyl-derivatives
2.3.1.9
1.1.1.36
(R)-3-Hydroxy-butanoyl-CoA
4.2.1.55
1.3.1.85
5.4.99.1
L-Glutamate
Alanine, aspartate and glutamate metabolism
1.1.1.93
5.1.2.5
1.1.1.37
Citrate cycle
Succinyl-CoA
(2S)-Ethyl-malonyl-CoA
4.3.1.2
CO2
Pyruvate metabolism
Formyl-phosphate
3.5.1.8
3.5.1.31
3.5.1.9
3.5.1.49
3.5.1.10
3.5.1.56
3.5.1.68
(S)-Malate
5.4.99.2
5.1.99.1
1.17.1.9
2.7.2.6
meso-Tartrate
trans-2,3-Epoxy-succinate
6.2.1.9
3.1.2.30
(R)-Methyl-malonyl-CoA
(2R)-Ethyl-malonyl-CoA
Mesaconate
1.2.2.1
cis-Aconitate
Oxaloglycolate
1.1.1.93
3.3.2.4
(S)-Malyl-CoA
2.3.3.9
5.4.99.63
Formate
5.1.99.1
2.8.3.-
3.1.2.10
Formyl-CoA
1.3.1.7
(S)-Methyl-malonyl-CoA
(2S)-Methyl-succinyl-CoA
1.2.3.4
Nitrogen metabolism
Ascorbate and aldarate metabolism
4.1.3.24
Acetyl-CoA
6.4.1.3
L-erythro-3-Methylmalyl-CoA
1.3.8.12
4.1.1.2
4.1.1.8
4.2.1.3
Propionyl-CoA
4.2.1.148
Oxalate
2.8.3.2
6.2.1.8
Oxalyl-CoA
4.1.3.24
Mesaconyl-CoA
1.2.3.5
1.2.1.17
Dihydroxy-fumarate
4.1.1.54
Tartronate semialdehyde
4.1.1.47
Glyoxylate
4.1.3.1
Isocitrate
Succinate
Glycolate
1.1.99.14
1.1.3.15
4.1.3.16
4.1.3.42
4-Hydroxy-2-oxoglutarate
Pyruvate
H2O2
O2
1.1.1.92
1.1.1.60
5.3.1.22
1.2.1.21
1.1.1.26
4.1.3.13
3-Oxalomalate
Oxaloacetate
Vitamin B6 metabolism
Purine metabolism
2.3.3.7
3-Ethylmalate
1.1.1.26
D-Glycerate
1.1.1.26
1.1.1.79
1.1.1.29
1.1.1.81
4.1.1.40
3.1.3.18
1.1.1.79
1.1.1.29
Butanoyl-CoA
Hydroxy-pyruvate
Glycolaldehyde
Phosphoglycolate
2.3.3.12
3-Propylmalate
2.7.1.165
2.7.1.31
1.1.1.77
Ethane-1,2-diol
D-Ribulose 1,5-P2
Pentanoyl-CoA
2-Phospho-D-glycerate
3-Phospho-D-glycerate
4.1.1.39
Carbon fixation
2.2.1.5
2-Hydroxy-3-oxoadipate
2-Oxoglutarate
Alanine, aspartate and glutamate metabolism
L-Glutamine
6.3.1.2
Carbon fixation
2-Oxoglutarate
1.4.7.1
L-Glutamate
Glycolysis / Gluconeogenesis
Glycine
NH3
GDC
2.6.1.4
2.1.2.1
2.6.1.45
L-Serine
Glycine, serine and threonine metabolism
00630 11/22/17
(c) Kanehisa Laboratories

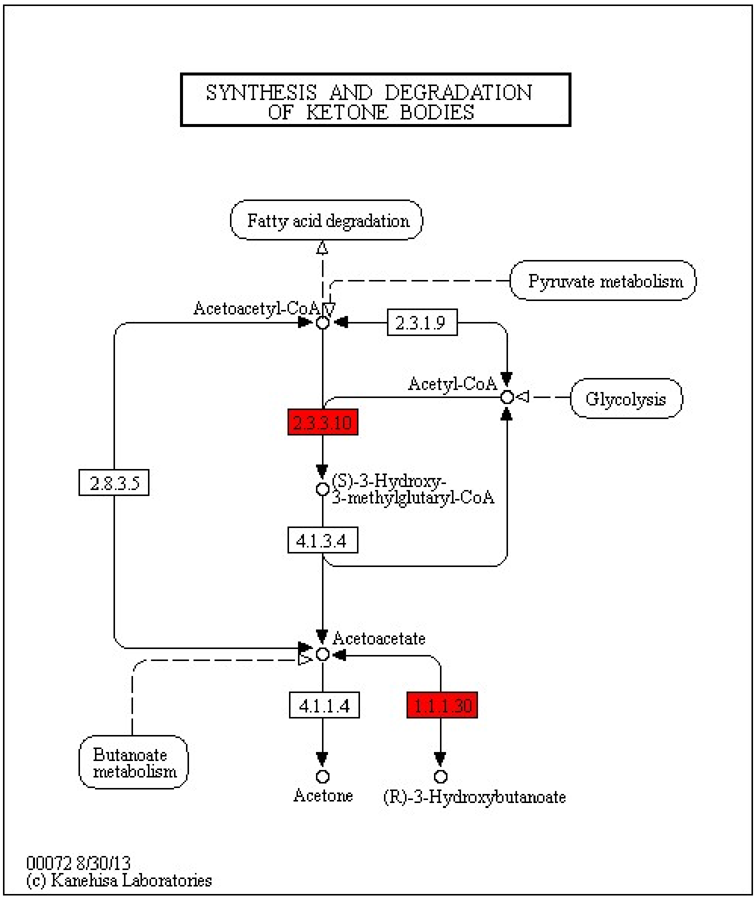
SYNTHESIS AND DEGRADATION OF KETONE BODIES
Fatty acid degradation
Pyruvate metabolism
Acetoacetyl-CoA
2.3.1.9
Acetyl-CoA
Glycolysis
2.3.3.10
2.8.3.5
(S)-3-Hydroxy-3-methylglutaryl-CoA
4.1.3.4
Acetoacetate
4.1.1.4
1.1.1.30
Butanoate metabolism
Acetone
(R)-3-Hydroxybutanoate
00072 8/30/13
(c) Kanehisa Laboratories

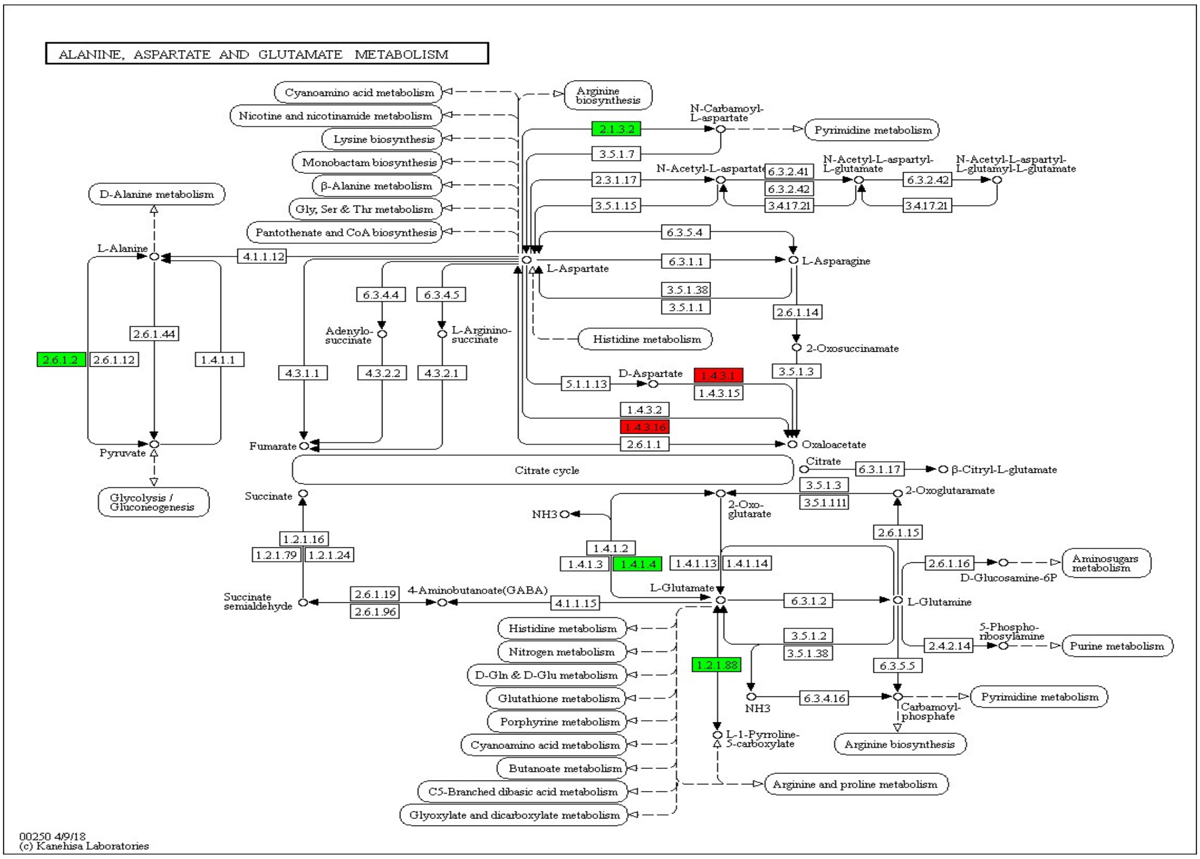
ALANINE, ASPARTATE AND GLUTAMATE METABOLISM
Cyanoamino acid metabolism
Nicotine and nicotinamide metabolism
Lysine biosynthesis
Monobactam biosynthesis
β-Alanine metabolism
Gly, Ser & Thr metabolism
Pantothenate and CoA biosynthesis
Arginine biosynthesis
N-Carbamoyl-L-aspartate
Pyrimidine metabolism
2.1.3.2
3.5.1.7
2.3.1.17
3.5.1.15
N-Acetyl-L-aspartate
6.3.2.41
6.3.2.42
3.4.17.21
N-Acetyl-L-aspartyl-L-glutamate
N-Acetyl-L-aspartyl-L-glutamyl-L-glutamate
D-Alanine metabolism
L-Alanine
4.1.1.12
6.3.5.4
6.3.1.1
L-Asparagine
L-Aspartate
3.5.1.38
3.5.1.1
2.6.1.14
2-Oxosuccinamate
6.3.4.4
6.3.4.5
Adenylo-succinate
L-Arginino-succinate
Histidine metabolism
2.6.1.44
2.6.1.2
2.6.1.12
1.4.1.1
4.3.1.1
4.3.2.2
4.3.2.1
D-Aspartate
5.1.1.13
1.4.3.1
1.4.3.15
3.5.1.3
1.4.3.2
1.4.3.16
2.6.1.1
Oxaloacetate
Fumarate
Pyruvate
Citrate cycle
Citrate
6.3.1.17
β-Citryl-L-glutamate
3.5.1.3
3.5.1.111
2-Oxoglutaramate
Glycolysis / Gluconeogenesis
Succinate
NH3
2-Oxo-glutarate
1.2.1.16
1.2.1.79
1.2.1.24
1.4.1.2
1.4.1.3
1.4.1.4
1.4.1.13
1.4.1.14
2.6.1.15
2.6.1.16
Aminosugars metabolism
D-Glucosamine-6P
Succinate semialdehyde
2.6.1.19
2.6.1.96
4-Aminobutanoate(GABA)
4.1.1.15
L-Glutamate
6.3.1.2
L-Glutamine
5-Phospho-ribosylamine
2.4.2.14
Purine metabolism
Histidine metabolism
Nitrogen metabolism
D-Gln & D-Glu metabolism
Glutathione metabolism
Porphyrine metabolism
Cyanoamino acid metabolism
Butanoate metabolism
C5-Branched dibasic acid metabolism
Glyoxylate and dicarboxylate metabolism
3.5.1.2
3.5.1.38
1.2.1.88
6.3.5.5
NH3
6.3.4.16
Carbamoyl-phosphate
Pyrimidine metabolism
L-1-Pyrroline-5-carboxylate
Arginine biosynthesis
Arginine and proline metabolism
00250 4/9/18
(c) Kanehisa Laboratories

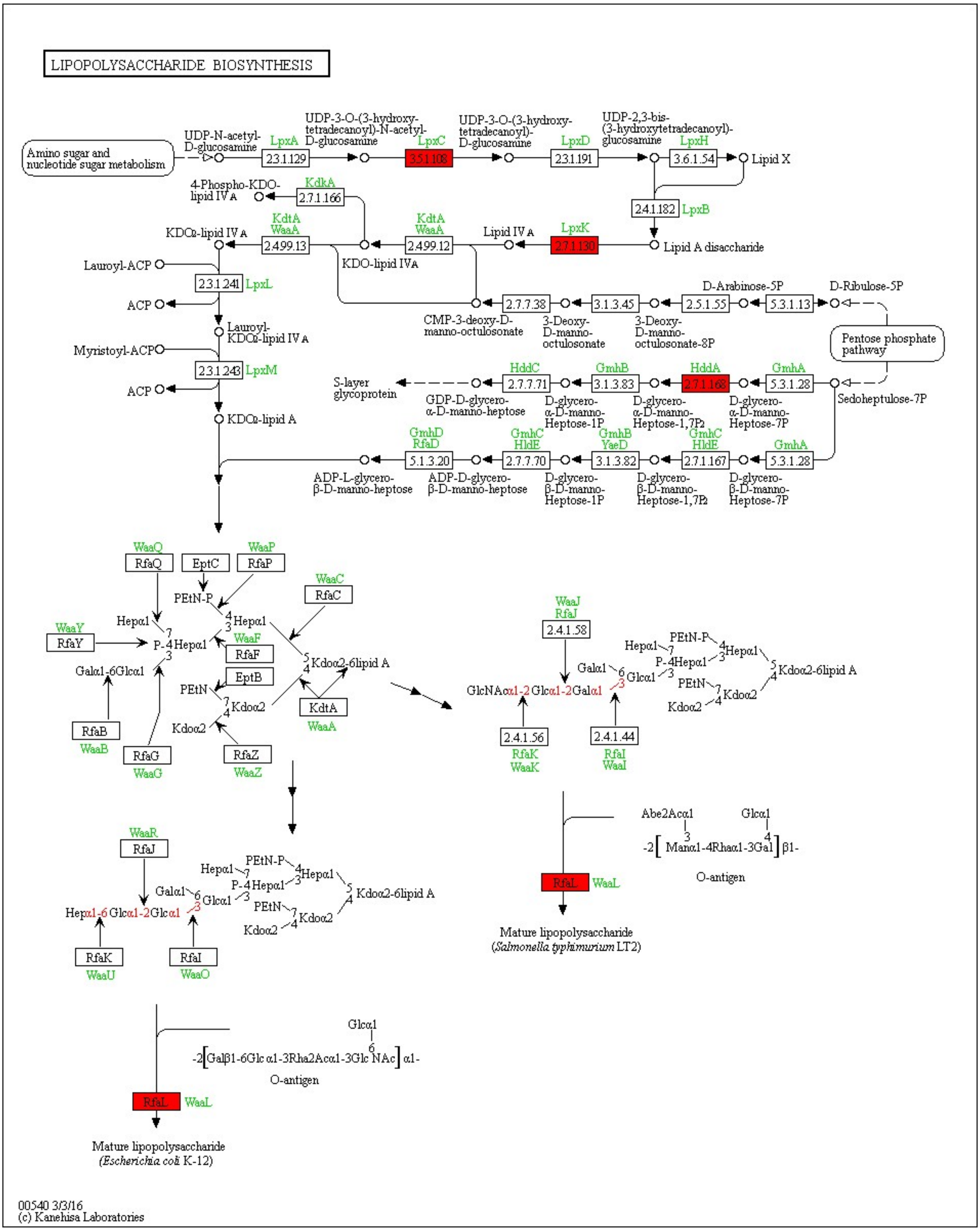
LIPOPOLYSACCHARIDE BIOSYNTHESIS
Amino sugar and nucleotide sugar metabolism
UDP-N-acetyl-D-glucosamine
LpxA
2.3.1.129
UDP-3-O-(3-hydroxy-tetradecanoyl)-N-acetyl-D-glucosamine
LpxC
3.5.1.108
UDP-3-O-(3-hydroxy-tetradecanoyl)-D-glucosamine
LpxD
2.3.1.191
UDP-2,3-bis-(3-hydroxytetradecanoyl)-glucosamine
LpxH
3.6.1.54
Lipid X
2.4.1.182
LpxB
Lipid A disaccharide
LpxK
2.7.1.130
Lipid IVA
KdtA
WaaA
2.4.99.12
KDO-lipid IVA
2.4.99.13
KDO2-lipid IVA
KdkA
2.7.1.166
4-Phospho-KDO-lipid IVA
Lauroyl-ACP
2.3.1.241
LpxL
ACP
Lauroyl-KDO2-lipid IVA
Myristoyl-ACP
2.3.1.243
LpxM
KDO2-lipid A
CMP-3-deoxy-D-manno-octulosonate
2.7.7.38
3-Deoxy-D-manno-octulosonate
3.1.3.45
3-Deoxy-D-manno-octulosonate-8P
2.5.1.55
D-Arabinose-5P
5.3.1.13
D-Ribulose-5P
Pentose phosphate pathway
Sedoheptulose-7P
GmhA
5.3.1.28
D-glycero-α-D-manno-Heptose-7P
HddA
2.7.1.168
D-glycero-α-D-manno-Heptose-1,7P2
GmhB
3.1.3.83
D-glycero-α-D-manno-Heptose-1P
HddC
2.7.7.71
GDP-D-glycero-α-D-manno-heptose
S-layer glycoprotein
D-glycero-β-D-manno-Heptose-7P
GmhC
HldE
2.7.1.167
D-glycero-β-D-manno-Heptose-1,7P2
GmhB
YaeD
3.1.3.82
D-glycero-β-D-manno-Heptose-1P
GmhC
HldE
2.7.7.70
ADP-D-glycero-β-D-manno-heptose
GmhD
RfaD
5.1.3.20
ADP-L-glycero-β-D-manno-heptose
WaaQ RfaQ
EptC
WaaP RfaP
WaaC RfaC
WaaY RfaY
WaaF RfaF
EptB
KdtA WaaA
RfaB WaaB
RfaG WaaG
RfaZ WaaZ
PEtN-P
Hepα1
P-4Hepα1
Galα1-6Glcα1
PEtN
Kdoα2
Kdoα2-6lipid A
WaaJ RfaJ
2.4.1.58
GlcNAcα1-2Glcα1-2Galα1
2.4.1.56 RfaK WaaK
2.4.1.44 RfaI WaaI
WaaR RfaJ
Hepα1-6Glcα1-2Glcα1
RfaK WaaU
RfaI WaaO
Abe2Acα1
Glcα1
-2[Manα1-4Rhaα1-3Gal]β1-
O-antigen
RfaL WaaL
Mature lipopolysaccharide (Salmonella typhimurium LT2)
-2[Galβ1-6Glcα1-3Rha2Acα1-3GlcNAc]α1-
O-antigen
RfaL WaaL
Mature lipopolysaccharide (Escherichia coli K-12)
00540 3/3/16
(c) Kanehisa Laboratories

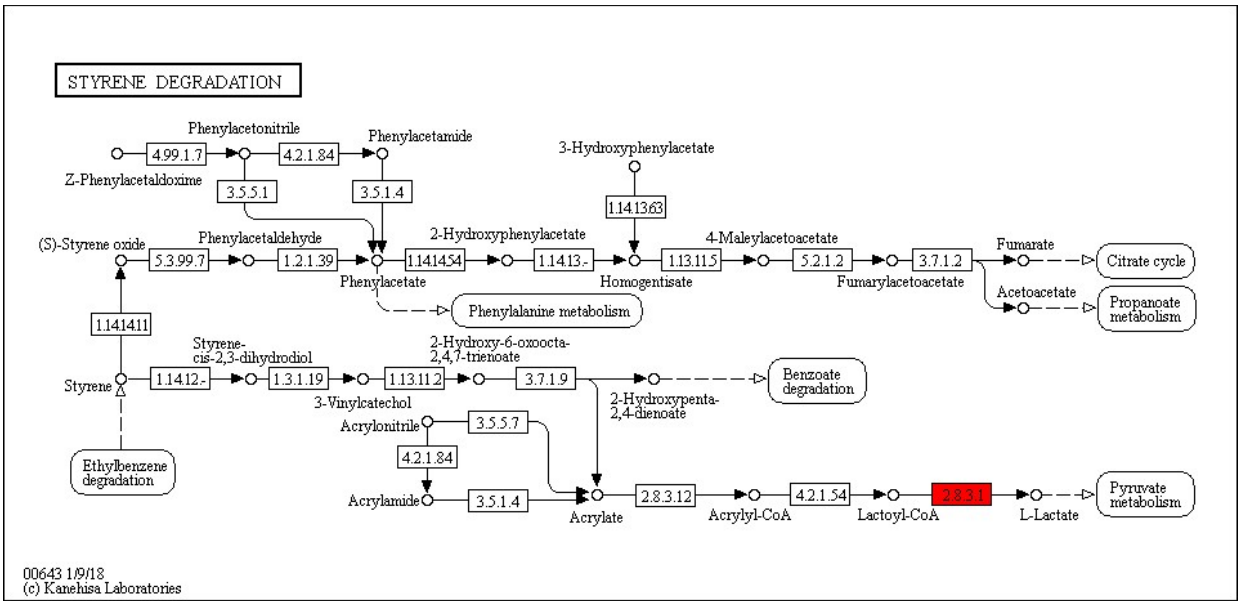


Fig.S5 Metatranscriptomic sequencing results of energy metabolism pathways (A) and KEGG metabolic pathway analysis (B)

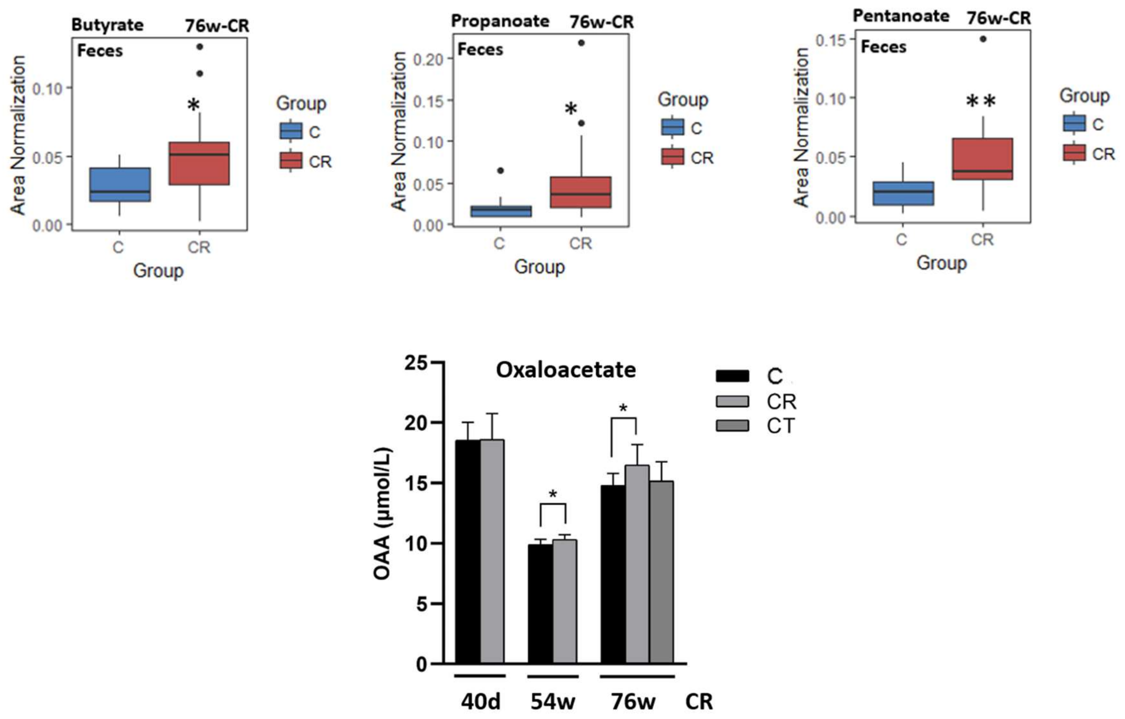


Fig. S6 Concentrations of propanoate, butyrate, and pentanoate in the rat feces in 76w-CR; the serum level of oxaloacetate in different CR experiments.

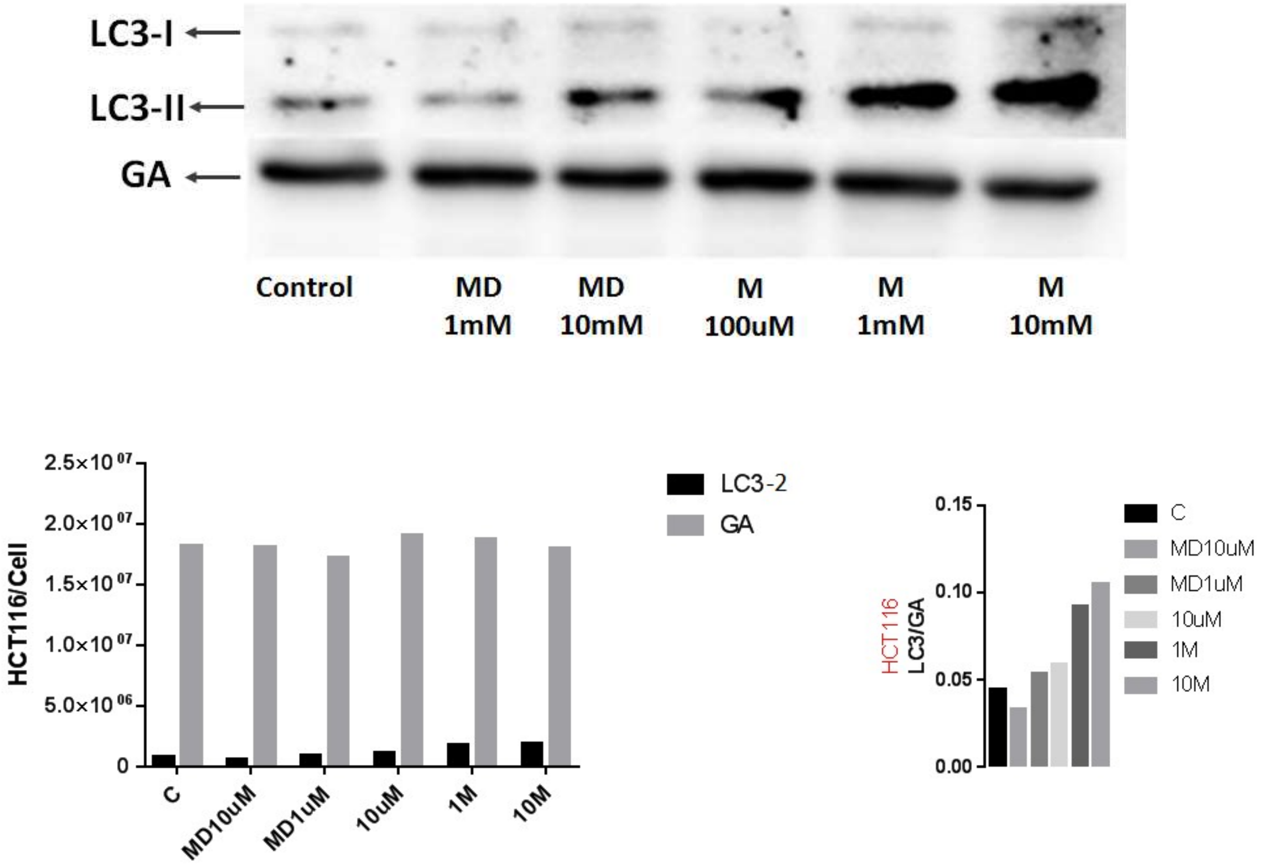


Fig. S7. Malate increased LC3-Ⅱin HCT116 cell line Sodium malate (M) had more effect than dimethyl malate (MD).

Table. S2 Differentially expressed transcription genes from Metatranscriptomic sequencing

| KO_L2 | KO_L3 | Definition | name.EC | Average C | Average CR | $FC_{(CR/C)}$ | P value |
|---|---|---|---|---|---|---|---|
| **Carbohydrate metabolism** | Pyruvate metabolism [PATH:ko00620] | pct | propionate CoA-transferase [EC:2.8.3.1] | 2734.600146 | 5811.9878 | 2.1254 | 0.0091 |
| **Carbohydrate metabolism** | Butanoate metabolism [PATH:ko00650] | E1.1.1.30, bdh | 3-hydroxybutyrate dehydrogenase [EC:1.1.1.30] | 11.461324 | 40.11462 | 3.5000 | 0.0008 |
| **Carbohydrate metabolism** | Butanoate metabolism [PATH:ko00650] | E2.3.3.10 | hydroxymethylglutaryl-CoA synthase [EC:2.3.3.10] | 47.761258 | 371.6416 | 7.7812 | 0.0019 |
| **Amino acid metabolism** | Alanine, aspartate and glutamate metabolism [PATH:ko00250] | nadB | L-aspartate oxidase [EC:1.4.3.16] | 85.71136 | 209.733144 | 2.4470 | 0.0020 |
| **Amino acid metabolism** | Alanine, aspartate and glutamate metabolism [PATH:ko00250] | DDO | D-aspartate oxidase [EC:1.4.3.1] | 6.757092 | 27.895398 | 4.1283 | 0.0366 |
| **Amino acid metabolism** | Alanine, aspartate and glutamate metabolism [PATH:ko00250] | E1.2.1.88 | 1-pyrroline-5-carboxylate dehydrogenase [EC:1.2.1.88] | 751.209404 | 318.642344 | 0.4242 | 0.0318 |
| **Amino acid metabolism** | Alanine, aspartate and glutamate metabolism [PATH:ko00250] | E1.4.1.4, gdhA | glutamate dehydrogenase (NADP+) [EC:1.4.1.4] | 18552.7194 | 5274.82372 | 0.2843 | 0.0182 |
| **Amino acid metabolism** | Alanine, aspartate and glutamate metabolism [PATH:ko00250] | GPT, ALT | alanine transaminase [EC:2.6.1.2] | 33.67002 | 11.447806 | 0.3400 | 0.0358 |
| **Carbohydrate metabolism** | Glyoxylate and dicarboxylate metabolism [PATH:ko00630] | DLD, lpd, pdhD | dihydrolipoamide dehydrogenase [EC:1.8.1.4] | 2840.554988 | 5929.1336 | 2.0873 | 0.0503 |

Table S3 and S4 are in the excel file.
