## Supplementary material for "Caloric Restriction Remodels Energy Metabolic Pathways of Gut Microbiota and Promotes Host Autophagy": SI-TableS3

**Supplementary Table S3. Environmental microbes' RNA sequences of isocitrate lyase and malate s**

**Sequences**

>CL48332Contig1
CTCGAACTGCTTGCGGTCGTGGAAAAGCTGCGCGCGAAACAACTGATCGACTTTGTTCGG
CACTGTGTCATATGGGTAATCTGCCACGTCTGGTCCTGGCTCGTTGGATGTGGACGCAGT
GGAACTGCACTGCCAGCCACTGACATACACCGAGGTCAAATACTTGGCCATCTGTGTCAC
CTGCACTGTGTCCAAGGCACCGAAGGTGTGGGAGCAGGTACCATTTGCCTGATGATCCCT
CATCATCTTGTACAGCTTGTCTGAAACCTTGGCGCCCGTGAAGTGCAGTGGCATGGTTCC
CTGCAGCTTCACAACATCCTCGACCCTGTAAGGCCTGGTGGTATGCTTGAAGCGTTCCTG
ACTCATCCATGCTGCCATGTCCTTGCATCGGCTTGTGAAGTCCGCATCCTCAGTCGAGCT
GGTGGATCGCATCTCGATA
>CL67599Contig1
GGGCTTCTGCTGGCAGTTCATCACACTTGCAGGCTTCCATTGTGATGCGCTGAGCATCGA
CCTCTTTGCCCGGGACTATGCGAAGCGCGGCGCGGCAGCCTATGTGCAGCTGATTCAGCG
AAAGGAGCGCGAGAACAAGGTGGAGACCTTGACACATCAGAAGTGGAGTGGCTCCGAGAT
CGTTGACGAGATGGGCAACATCATCTCAGGAGGAACTTCTTCGACAGGAATCATGAGTGC
TGGTGTGACCGAGAAGCAGTTCTAATGTTTGCTATGCCACGATTTTACCGCAAGGTAGGT
TTCGTTGCTTCATTTTCAAAATGTGAAGTTGTTTCGATTTTCTGTTTTCAGAGCAAGTAG
TGTGGGCAAGGCTTCACATCGACCTATTTGCTTGCAAAGTCTCTTGACAGTTCCTTTGGA
CCGTGGCCGGACGAGCCCGTGAATGG
>CL125104Contig1
AGTTGTGACACTTCAGGGCACGCAACCTCTGTATTTTCACAGCGCCAAAATTGCAGAGAA
GCTCTACAACATGTCCCGTGAGCATCAGAAAAGCGGCACATGCTCTCACACATTTGGTGC
CCTCGATCCCGTGCAAGTTGTGCAGATGGCTCCGTACCTGAGCTCAGTATACATCTCGGG
ATGGCAAAGCTCATCCACCGCAAGCACCTCCAACGAGCCTGGTCCAGATGTTGCAGACTA
TCCATATGACACTGTGCCGAACAAGTGTGACCAGCTCTTCCGAGCACAGCTGTTCCATGA
C
>CL125491Contig1
GCCCGCAGCTCCGTTCTCGATGAACATCTTCGCAAGCTTCATGGTTGCGGTCAGGCCACC
ATGTCCGGTGTCAGCGTCCGCGATGATCGGGTTCAGGTAGTCCACGCAGGGCTGCTTGGA
GCGCTCCTCCGGCGACATGCGCCGGCGCTCCTCATACTGCTTGCGGTCGTGGAAGAGCTG
CGCGCGGAAGAGCTGGTCCACCTTGTTCGGCACGGTGTCGTAGGGGTAGTCGGCAACGTC
CGGGCCAGGCTCGTTGGAGGTGGAGGCAGTGGAGCTTGACTGCCAGCCACTGACGTAGAC
GCTGGTCAGGTACTTGGCCATCTGGACTACCTGCACCGTGTCCAGGGCGCCGAAGGTGTG
CGAGCAGGTTCCTTTTGCC
>CL1Contig2463
ACACGACGCTCTTCCGATCTTCTCCTTTGCTCGTCAAAGTCATCATCAAGAAAATGTGCG
GATGTGCACGTCAGAGGATGCGAAAATCGAAGGGCCCATTTGCTAGTGGGCATGCGCCTC
TGTCCGAAAATCTTAGAGCTTGTTCCGTTCTGAGAGAGCAAGCTTAAGAAGCTGTCGCTG
TTGACAACTCGTCTATGTGATCTGCGTTCAGAAAAGAAAAAGAGCCGCCTAATGCTTGCC
AAACTGAGACTCTGTTACGCCGGCGCTCATGATTCCTGTGGAGGCCTTGCCCCCAGTAAC
AACGTTCAGATATGCATCAACAATCTCAGCTCCAGACCACTTCTGATGAGTCAAGGTCTC
CACATCCTTATCCCGCTCTTCACGCTGAATCATTGTTACATAAGCTTTCATACCATCTGT
TTTGTATGCACGAGCGAACAATGAAATACCAAGGGAATCCATGTGGAAACCCGCAAGTGT
GATGAACTGCCAACAAAAACCCATCTTTGCAAGATCCCAAATGAA
>c664616_g1_i1
GCTCTTCACGCTGGATCATACTAACATAAGCGCCTGCCCCCTTTGTCGCATAGGCCTTTG
CGAAGTTGTCGATAGAAAGCGCATTGCAATGGAAGCCAGCGAGAGTGATGAACTGCCAGC
AGTAGCCGAGCTTGGCCAAGTCAAAAATGAAGGTCTCCATCTGCGCATCTGTCATGCCTG
AGTTGTCCCAGTTGAAGGAAGGGCTGTTGTTGTATGAAAGCATCTGGTGTGGTGCAGCTG
CCCTGACTTCCTTTGCGAATTGTGTTGCCTGGCTCAGGATGGGCTTGCCTT
>c892671_g1_i2
GCCTTCAGTGCTTTCGCCACTGCGTCTGGGAATGTCATGGCGCCAGCCCTCTCTTCCCAG

TCCTTGTCAGTACCTTTCTTGATCGCTTCATAAAGCGGCTCCACGCCTTTGACCGTCGCA
CCCTTGATGTGAGGGTGGTCGATTGGATCAATATTGGAATCAATCATGGTGGCAGCCTCT
GCATCGGTGCGGGCAACCAGAACAAGGGGTGAATTGAGGACGTCTGCCTGCAGCCGGATA
GCGATGAGGCGCTGGACGTGCTCCTGAGTAGACACAAGGACTTTGCCTCCCATGTGGCCA
CACTTCTTGGTTCCAGGCTTTTGGTCCTCGATGTGAATACCAGCTGAGCCATTTTCGACG
AACATTTTGGCAAGC
>c914354_g3_i1
CGCCGACTGCATCTGGATGGAGACGGGCAAGCCCATCCTCAGCCAGGCAACCAAGTTCGC
GAAGGAGGTGCGTGCGGCTGTGCCCCACCAGATGTTGGCCTATAACTTGAGCCCCTCCTT
CAATTGGGACTCCGCCGGCATGACGGACGCGCAGATGGAGAGCTTCATTTGGGACCTGGC
CAAGCTGGGCTTCGTGTGGCAGTTCATCACGCTTGCGGGGTTCCACTCCGACGCGCTCGG
CATCGACCTCTTCGCGAGGGAGTACGCCACGAAGGGCGCCGCCGCGTACGTGCAGCTCAT
CCAGCGGCAGGAGCGGGAGAAGGGCGTGGAGACGCTCACGCACCAGAAGTGGAGCGGCTC
GGAGATCGTGGACGAGGCGGGCAACATCATCTCCGGCGGCACCTCCTCCACAGGTATCAT
GAGCGCCGGCGTAACGGAGGCGCAGTTCGGCGCGAAGCATTGAGCGAGGAACGGCTGCAG
GAAACAGGCGAGTGGCGTCGCTTGCCGACAGGTGCTCGCGCCGCCCCGTCGCGCCCGCGT
CGCGGAGCCGAGGCGGTCTCGCCATGCCCCGGTCAGGATTCGAACGAAC
>c914354_g3_i2
GGCAAGCCGATCTTGAGCCAGGCCGAGAAGTTTGCGAAGGAGGTACGCGCAGCAATCCCG
CACCAGATGTTGGCGTACAACTTGAGCCCTTCCTTCAATTGGGACTCGTCCGGCATGACG
GACGCGCAGATGGAGAGCTTCATTTGGGACCTGGCCAAGCTGGGCTTCGTGTGGCAGTTC
ATCACGCTTGCGGGGTTCCACTCCGACGCGCTCGGCATCGACCTCTTCGCGAGGGAGTAC
GCCACGAAGGGCGCCGCC
>c914354_g3_i3
CGAAGGAGGTGCGCGCCGCGGTGCCCCACCAGATGCTCGCGTACAACCTGAGCCCCTCCT
TCAACTGGGACTCGGCCGGCATGACGGACGCGCAGATGGAGAGCTTCATTTGGGACCTGG
CCAAGCTGGGCTTCGTGTGGCAGTTCATCACGCTTGCGGGGTTCCACTCCGACGCGCTCG
GCATCGACCTCTTCGCGAGGGAGTACGCGACGCGTGGCGCGGCGGCCTACGTGCAGCTCA
TCCAGCGCAAGGAGCGCGAGAACGGCGTGGAGACCCTGACGCATCAGAAGTGGAGCGGCT
CCGAGATC
>c909308_g1_i1
ATTTTCAATGAACATCTTTGCCAGTTTCATCGTTGCTGTGAGCCCTCCGTGACCAGTGTC
GGCATCGGCAATGATTGGGTTTAAGTAGTCTACAAGTGCCTTTTTGCTTCGCTCCTCAGG
GCTCATGCGTCGTCTCTCCTCATACTGTTTACGATCATGAAAGAGCTGCGCTCGAAACAA
CTGGTCCACTTTGTTTGGAACCGTATCATACGGG
>c692422_g1_i1
CAGAAAAAGAACAGCTTAAAGAAGTGGATGAGTTCTTCAATGACCCTCGCTTCAAACACA
CGACTCGGGGTTATGACGCTGGCACAGTCGCTCGTCTTCGTAGCCCAATTGAACGGAATT
ATGCGGGTACAGCACCAGCAGAAAAGTTCCATTGTATGATTCAAGACCTTGCCAAAAAGG
GGGGGTTTAGTCACACTTTTGGGTGCTTGGATCCAGTTCAGGTGATCCAAATGGCTCCCT
ACCTCACCTCGATTTACGTGAGTGGCTGGCAATCCTCTTCGACTGCCTCTACCACCAACG
AGCCGGGGCCCGATCTTGCTGACTACCCCTACACAACAG
>CL1Contig10513
CGCATTTCTTTGTGCCTGGACTCTGATCCTCGATGTGAATGCCAGACGCACCAGCCTCGC
AAAACATTTTCGTCAACTTCATAGTGGCAGTCAAACCGCCGTGTCCAGTATCAGCATCCG
CGATGATTGGATTCAGAAAATCGGTTGCTGGATGTTTGGCCCTCCATTCGGGCGACTCCC
GCATCCGAGCTTCTTGTTGCTTGCGATCGTGAAAATGCTGAGCCTTGAACAACTGTTCGA
CCTTATTTGGTACAGTGTCCATCGGATAGTCTGCCACATCAGGCCCAGGCTCGTTGGAAG
TGCTTGCAGTGCTGCTACATTGCCAACCACTTACATATACACTCGTCAAGCCAGCTTCAG
CCATTTGAACTACTTGCACTGTAGACAATGCTCCGAAAGTGTGAGAGCATGTCCCAT
>CL1Contig12374
GCGTCTTGGCCTTTTGCCTTCAGTGCTTTAGCCACTGCGTCGGGGAATGTCATGGCGCCA
GCCCTCTCTTCCCAGTCCTTGTCAGTACCTTTCTTGATCGCTTCATAAAGCGGCTCCACG
CCTTTGACCGTCGCACCCTTGATGTGAGGGTGGTCGATTGGGTCGATGTTCGAATCGATC

ATGGTAGCAGCCTCTGCATCGGTGCGGGCAACCAAGACCAGGGGCGAATTGAGAACGTCA
GACTGCAGCCGAATGGCGATGAGACGCTGGACATGCTCCTGGGTAGAGACGAGGACTTTA
CCTCCCATGTGGCCGCACTTCTTGGTTCCAGGTTTCTGGTCCTCAATGTGAATACCAGCT
GCGCCGTTCTCAATGAACATCTTAGTAAGCTTCATCGTGGCAGTGATACCTCCGTGGCCT
GTGTCTGCGTCCGCGATGATGGGGTTAAAGTAATCGACAGCCGGAGTATTTGCTCTCTCA
TCAGGCTTCAAATGGCGGCGCTCCTCGAACTGTTTGCGGTCATGGAAAAGCTGTGCACGG
AAAAGCTGGTCCACCTTGTTGGGCACTGTGTCGTAGGGGTAGTCTGCCACGTCTGGGCCG
GGCTCATTGGAAGTGGAGGCCGTCGAGGAGCATTGCCATCCGCTGACATATACGGAAGTG
AGGTACTTTGCCATCTGGACCACTTGGACAGTGTCCAAAGCCCCGAAGGTGTGAGAGCAA
GTGCCCTTGGCCTGATGCTCG
>c775297_g1_i1
GGTGGAAGAGGTGGAGCTTTTGAAGCACCAGAAATGGAGTGGAGCTGAACTTGTTGATCG
TATGGTGAATGTGGCTAGTGGTGGACAATCCTCCACTGCTGCAATGGGAGCTGGTGTCAC
CGAGGATCAGTTTTCCAAGCACTGATTGATCAAATTTTGCACATGTAGTTGTTGAGCAGT
CTTGTCTTGTTTTATATCATCTCCTGGTGCTAAGTACAATAGGATACTCAGCTTATCTTA
ATCAATCTTCTCGCCTCCTAAATAGTTAAGACTGTGCTAGTACACTACCAAATGTAC
>c883116_g1_i1
GGTGGAGCTGCACTGCCAGCCGGAGACATACACGGAAGTCAAGTACTTGGCCATCTGCGT
CACCTGCACTGTGTCCAAAGCTCCGAAGGTGTGAGAGCAGGTGCCATTCGCTTGGTGCTC
CCGCATCATCTTGTAGAGCTTCTCCGACACCTTCGCACCGGTGAAGTGCAACGGCATGGT
GCCCTGAAGCTTCACCACATCCTCCACCTTGTACGGGCGTGTCGTGTGCTTGAAACGCTC
TGAAGACATCCAAGCTGCCATGTCCTTGCAGCGCTGGGCGAAATCAGCATCCTCGCTGCC
GCTCGTGGCCTTCATCTCCACACCACCAATCACGGCGGCATCCCCCGCCAAGACATGGCG
CTGCACTGCGAGAACACGATTCGCCATATTGCTAGCACGAGAACCTATTATTC
>c849717_g1_i1
CTGGTGATTCGCGGTCTCTGCCACATTCTCATCACCTTCTAGCTGGATGAACGGGAAAAC
TTATTGAAAACAAAGATGCACTGAACAGGGGCAAATTGGCCTTCTGTGGTGAGCTTCTGT
AGCAAGCTCACAGGTGCATCCTACCATCTCTGTCAACCTCAGAACTGCTTCTCTGTAACG
CCTGCAGACATGATGCCAGTTGACGAGAGGCCGCCTGAGACAATGTTTCCCATCTCGTCC
ACGATCTCGGAGCCACTCCACTTCTGATGAGTCAGCGTCTCGACTTTGTTTTCACGCTCC
TTCCGCTGGATCAGCTGCACGTAAGCCGCAGCTCCCCGTTTTGCATAGTCCCGCGCGAAG
AGATCGATGCTCAGGGCGTCGCAGTGGAAGCCAGCAAGTGTGATAAACTGCCAGCAAAAG
CCAAGCTTGGCAAGGTCCCAAATAAAAGACTCCATCTGGGCATCAGTCATCCCAGCTGTG
TCCCAATTGAAGGAAGGGCTCAGGTTGTAG
>c902440_g2_i1
GACGCCGGCACTCATGATCCCCGTGGACGACAACCCGCCAGAGATGATGTTCCCGGCTTC
GTCGACAAGCTCCGATCCGCTCCACTTCTGGTGCGTCAGCGTCTCCACTCCCTCTTCTCG
CTCCCTGCGCTGAATGAGCTGCACGTACGCCGCCGCGCCCCGCTTGGCGTAGTCGCGCGC
GAAGAGGTCGATGCTCAGGGCGTCGCAATGGAACCCTGCCAAGGTAAT
>c738735_g1_i1
GCCGAATTGATCCTCCGTCACACCTGCTCCCATGGCTGCAGTAGATGATTGCCCACCGCT
TGCTACATTAACCATCTTATCCACAAGTTCTGCACCACTCCATTTCTGATGCTTCAATAG
CTCCACTTCTTCAACCTTCTCTTGCCTCTGAATGTCCCTAACATAAGCAAGCATGCCCTC
ATCTCCAAATGATCGTGCCAACTTTGTTACAATGAGACCATTCGAGTGAAACCCAGCCAA
GGTAATGAATTGCCATGCATACCCAAGCTTCCCAAGATCATCAT
>c810828_g1_i1
GACGCTCTTCCGATCTCATCCTCACATCAAGGGTGCGACGAAGCAGGGTATCGAGCCCCT
GTACGAGGCCATTCGAAAGGGCACCGACAAGGGCTGGGAGGAGCGGGCGGGCTGCATGAC
CTTCCCGGATGCTGTGGCCGCCCAGTTGAAGGCGAAGGGCAAGGATGCCAGCCAGTGGCT
GAAGGATTCTTTGAAGATGTCCCTCGACCAGATGCGGAAGGCAGCCAAGGACATGGGTTG
TGAGGTGTTCTTTGACTGGGATGCCGCTCGGTCTGTGGAGGGTTACTTCAGGATCAAAGG
CTCCACGGAGTTTTGCATCCAGCGTGCCATCGCTTGGGCACCGTATGCGGACTGCATCTG
GATGGAGACCGGC
>c892052_g1_i1

CCTCACCAGATGCTGGCCTACAACCTGAGCCCTTCCTTCAACTGGGACACGGCTGGGATG
ACGGATGCCCAGATGGAGTCCTTTATTTGGGATCTTGCCAAACTTGGTTTCTGTTGGCAA
TTCATCACGCTTGCGGGCTTCCACTGCGATGCCTTGAGCATTGATCTTTTCGCACGGGAT
TACGCCAAGCGGGGCGCTGCAGCTTATGTGCAGCTCATCCAGCGAAAGGAGCGCGAGAAC
AAGGTGGAGACATTAACTCACCAGAAATGGAGCGGCTCCGAGATCGTAGACGAGATGGGG
AACATAGTGTCAGGAGGCATGTCTTCCACTGGAATCATGTCGGCGGGCGTCACAGAGAAA
CAGTTCTGAGGCGCGCTGGGTTGCAAAACTGTGAGGTACGCAGCATGTCCAAGGACGTGA
GGTTTGACGCCGCTAGTTGTTTCGTTTTTCTCAGTTTTCCTCCGGTGAAGCTGAAGGGTT
AACAACGGGG
>c810024_g1_i1
ATCCATTCATTCTTGGAGCAACCACTCCTGGCTCTATCTCGCTAAGTGAAGCATTGGCAG
CTGCAGCAGCCAGTGGAGGGGCAAATGCAGCTGATGTGGAGAAGGAATGGACTGCAAAAG
CTCGTCCAATGACTTTCAGTGAGGCAGTATTAGATAAGATTCATTCATTGCATGTAGATG
GAGGAAGAAAGAGAGAAATGGTCAAGATGTGGCAAGCGAGTGATCCTGATGCATTAAGCA
ACGCAGCTGCCCATCGCGTAGCAGACTCTATTTTTGGGAAGAAGAACTCCATATACTTAG
ACTGGGAAGCATGTCGTGTTCGTGAAGGATACTATCGTATTAAACCAGGG
>c398051_g1_i1
GTGGAAACTTTGACACACCAGAAGTGGAGTGGGTCAGAACTCGTGGATGCATATTTGAAT
ACAGTCACTGGTGGGACTGCTTCGATTTCCATCATGGGTGCTGGCGTGACTGAGACACAA
TTTGGTGCCAAGTCTAAGCTTTGATTGACCGCTTCCAGACAAGCTGCTTCACTTTACTGG
CATTGGCCAAGCAGTGGGGCAACTTTTTGAGTTTCGAGTATTTTCGTTCCAAGCGAGCAA
TTTTCGTGCCGAG
>c810371_g3_i4
AGGGGGTGAAAAATGAGATGCGAGATAAGCTGGTCTCACGCGATGGCCGGGCGCCAGGTT
CAGCTACGACCTATGCCTATTCGTGAAGCCTGGCCTAAAGTCGGACACCTCTCCGTGTGA
GGGAGAATTACTTGGCGCCGAATTGGGTCTCGGTCACGCCGGCGCTCATGATGCCGGTGG
ACGAGGTGCCGCCAGAGATGATGTTGCCCATCTCGTCCACGATCTCGGAGCCGCTCCACT
TCTGGTGGGTCAGGGTCTCCACCTTCTCCTCGCGCTCCTTCCGCTGGATCATCTGCACGT
AGGCCGCGGCGCCCTTCGTTGCGTATGAGCGCGCGAAGAGGTCGATGCTGAGGGCGTCGC
AGTGGAACCCTGCCAGGGTGATGAACTGCCAGCAGAAGCCGAGCTTGGCGAGGTCCCAGA
TGAAGGTC
>c524822_g1_i1
GCCCTCACCAACACGCAAAGAACGAATCTTATCCAATACTGCCTCTCCAAATGTCATTGG
CCGAGCTTTGGCAGTCCATTCCTTTTCGATACTTGCTGCATCTCCTCCTCCAGATGCAGC
TGCATTGGCAACAGCCTCTCTCATTGATACAGATCCAGGCACGGTTGCTCCCAAAATGAA
GGGATGATCTCTTCCATCACAGTTCGAATCTAACAATGTAGCTGCCTCAGCATCAGTCCT
TGCCACTATGACCAACTCCACGCCTAATACATCTGCAGCCAAACGTGCTGCAATCAATCG
ATCTGCGTGCTCTTGCGTGGCAACTAAAACTTTACCTCCCATATGGCCGCACTTTTTCGT
GCCAGGCTTTTGATCTTCGAAGTGGACCCCAGCAGCTCCAGCCTCTACAAAGAGTTTCAC
AAGTTTCATAACAGCACTTAAGCCTCCATGTCCAGTATCGCCATCGGCAACGACGGGTGT
AAGATAATCGACCTTTGGTCCTGGGTTTTGACCATTTAGGATCGCAGATGCACGTTCTTC
ATTCTGACGACGATCATGATGAAGCTGGGCTCTCACAAGTTGATCACATTTCTT
>c736104_g1_i1
AGAAGGGTTTTATAGGTTCCAAGGTTCCACCAAGGCTGCTATTCTCAGGGGTTGGGCTTT
TGGCCCCCATGCAGACCTAATTTGGATGGAGACATCAAGCCCCGACATGGTAGAATGTAA
GGAGTTTGCAGAAGGGGTGAAATCAAAGCATCCAGACCTGATGTTGGCCTACAATCTGTC
ACCTTCTTTCAATTGGGATGCTTCTCGCATGAGCGATGAACAGATGGTGAACTTCATTCC
TCAG
>c836096_g1_i1
CTCCGCATGTCACCAGAGGAGCGTTCAAAGACACCTGTAGTGGACTATTTGAATCCAATC
ATTGCTGATGCCGACACGGGCCATGGAGGCCTCACAGCCACCATGAAGCTTGCGAAGATG
TTTGTCGAAAACGGCGCAGCAGGCATTCACATTGAGGACCAGAAGCCAGGTACCAAGAAG
TGCGGACACATGGGTGGCAAAGTCTTAGTTTCGACACAGGAGCACATCCAGCGGCTTATT
GCCATCCGTTTGCAGGCAGATGTCCTCAACTCTCCTCTCGTGGTCGTTGCCCGCACAGAT

GCGGAAGCTGCCACTATGATCGATTCCAACATCGACTCCATCGACCATCCTCACATCAAG
GGTGCCACAGTGAAGGGAGTCGAGCCACTCTACGAGGCCATCAAGAAAGGAACTGACAAA
GACTGGGAGCAG
>c736768_g2_i1
GGGGGTTTGCGATTCTTTCTTCCGCAGTCATCTGTGACCTTTCAAAGCGCTGTCGTCTGT
CGTGGAAGCATTGTGCCTTGAACAGCTGTTCTACTTTGTTTGGAACGGTATTAGCAGGAT
AGTCAGCAAAGTCTGGACCGGGCTCATTGGAACTGGAGGCGGTGCTACTGCATTGCCAAC
CACTGACGTAGACCGTTTCGAGATATTTGGCCATTTGCGTCACTTGTACGGT
>c892052_g1_i2
CCTCACCAGATGCTGGCCTACAACCTGAGCCCTTCCTTCAACTGGGACACGGCTGGGATG
ACGGATGCCCAGATGGAGTCCTTTATTTGGGATCTTGCCAAGCTTGGCTTCTGCTGGCAG
TTCATCACACTTGCAGGCTTCCATTGTGATGCACTGAGCATCGACCTGTTTGCCCGGGAC
TATGCGAAGCGTGGTGCGGCAGCCTACGTGCAGCTCATTCAGCGAAAGGAGCGCGAAAAC
AAGGTTGAGACTTTGACACATCAGAAGTGGAGTG
>c907594_g1_i3
CGGCCACATGGGCGGCAAGGTGCTTGTGAGCACCTGGGAGCACATCCAGCGCTTGATCGC
CTGCCGGCTGCAGGCCGACGTGCTCAACTCGCCGCTGGTGCTGGTGGCGCGCACCGACGC
AGAGGCCGCGACCATGATCGACTCCAACATCGACCCCATCGACCACCCCCACATCAAGGG
AGCCACCGTGCAGGGCGTGGAGACTCTGTACGATGCCATCAGGAAGGGCACTGACAAGGA
CTGGGAGGAGCGCGCCGGCTGCATG
>c357732_g1_i1
GAGGCCTGCCGTGTGCGCGAAGGTTACTATCGAATTAAGCCCGGGGTCGAATACTGCATC
CAACGTGCCCGTGCCTACGCTCCTCATTCAGACTTGATCTGGATGGAAACAGGCAAGCCG
GGAATTCCAGTTGCTCGCAAGTTCAGCCAAGGTGTAAAGGCTGTATTTCCACACCAAATG
CTTGCGTATAACCTGAGCCCATCATTTAATTGGGATGCGAGTGGCATGTCAGATGAAGAA
CTTGCAAATTTCAATGATGATCTAGGAAAACTTGGATATGTTTGGCAATTCATAACACTC
GCGGGGTTTCATTCAAATGGTCTTATTGTCACAAAACTTGCTCGTTCTTTTGGAGACGAG
GGAATGCTGGCATATGTCCGAGATATTCAAAGACAAGAAAAGGTGC
>c933242_g1_i1
GGGGTATTACATGATCAAGCATGGTGAGCAGTATGCAATTGCACGCGGCCGTGCATTTGC
ATACCATGCGGACATCCTGTGGATGGAAACCGGAAAGCCTCTCCTAGCTCAGGCAGAAAT
TTTTGCAAAAGGCATCAAAGCTCTCCATCCAACACAGATGCTGGCGTACAATCTGTCACC
ATCATTTAACTGGGACGCTGCTGGAATGACCGATGAGGCAATGAAAAGTTTCATTTGGGA
TCTTGGTCGTCTTGGTTTCTGCTGGCAATTCATCACGCTTGCTGGTTTCCACACAAATGC
TCTGGGCCTTACAAATTTCGCAAGGGGTTACAGCAAGGAAGGGATGCTCTCGTATGTGCG
AGATGTGCAACGGAAGG
>c905333_g1_i2
CCTGAAGAGCTGGTCCACCTTGTTGGGCACGGTGTCGTAGGGGTAGTCCGCCACGTCGGG
GCCTGGCTCGTTGGAGGTCGACGCCGTGGAGCTGCACTGCCAACCGCTGACGTAGACGGA
GGTCAAGTACTTGGCCATCTGAACGACCTGGACGGTGTCCAAGGCGCCGAAAGTGTGGGA
GCAAGTGCCCTTGGCCTGGTGCTCACGGAGCATGGTGTAGAGCTTCTCGGAGACCTTGGC
ACCCGTGAAGTGCAACGGCAAAGTGCCCTGCAACTTCACGACGTCCTCAACTCTGTAGGG
CCGCGTAGTATGCTT
>c831242_g1_i1
GAAAAAGTGCGGCCACATGGCTGGCAAAGTGCTTGTTAGCGCCCAGGAGCACTGCAACCG
TTTGATTGCTGCCCGTCTGCAAGCCGACATCTTGGGCACTGAAACCCTAGTTGTTGCCCG
TACTGATGGTGAGGCCGCCACTCTGCTCGACAGCAACATCGACCCCCGCGACCACCCCTT
CATCCTGGGCTGCACCAACGAGAGCCTGCCCTCGCGTAACGATGCAATCCGCGGAAAGAA
GGGCGCCGCCTTCAGCGCCGCCGGCGCTGAATGGGACAAGCAGGCCAATCTCATGACCTA
CGGTGATTGCGTGGCCGCGGCTCTCAAGGCGGCCGGCAAGCCCACCGAGGAGTGGTTGGC
TAAAAGCGCCAAGATGGGCCACGATGAGGCACGCGCCTATGCCGCCTCCCTGGGCGTCAA
TCCGTACTGGTGCTGGGATAAGCCCCGCGCTGTGGAGGGCTACTTCAAGATCAAATGCGG
CGCTGACTACTGCATCGCACGTGCCATCGCCTACTCGCCATATGCCGACCTCATCTGGAT
GGAGACCGGCAAGCCTATTCTGGCACAGGCCACTCAGTTTGCCAAGGAGGTGCGCGCGGC

CGTGCCGCACCAGATGTTGTCTTACAACTTGAGCCCGTCCTTCAATTGGGACACAGCAGG
CATGACGGATGCACAGATGGAGTCCTTCATTTGGGACCTTGCGAAGTTGGGCTTCTGCTG
GCAGTTCATCA
>CL12084Contig2
CCACATCAGATGCTCGCGTACAACTTGAGCCCTTCATTCAATTGGGACACTTCAGGCATG
ACAGATGGTGAGATGGAAACCTTCATCTGGGATCTGGCAAAGATGGGATTCTGCTGGCAG
TTCATCACGCTCGCAGGATTCCATATGGATTCCCTTGGCATTTCGCTTTTCTCACGTGCA
TACAAAACCGATGGTATGAAGGCCTACGTAACAATGATTCAGCGCGAGGAGCGTGACAAG
GATGTGGAGACCTTGACCCATCAAAAATGGTCTGGAGCTGAGATCGTCGATGCTTATCTA
AATGTGGTAACTGGCGGCAAAGCATCTACAGGGATCATGAGTGCTGGTGTAACGGAGTCG
CAGTTTGGCAAGTGATAGTCGTGTTAGGCGTTTCGTTTTTCTTGTGCACTCAAGTTTTAT
AATCGTTGCTCGCGTGGCCATTGCTACAGTATTTTCGTGTTGAGCGCATAAACGCTCGCA
CTATGGCGAGCGAGCCATTTGCTTTTTCAGATTTCGGCCTGCAGCAAAGCTGGAAACTCG
GACACATGACTGAATAAGAGTAGCGCATCAGAGAAAAA
>CL53156Contig1
GTCCATCAGAAAACAACAACAAACAGTCCGCTGGAGGCCTGATTGATGTCAATCCACAAC
CAAACGGCCAATGCCGCGTGTGGGTACAGCAAACTTCATCAGCGAAGGCTTCGTCAAGGA
AAGTATCAAAGACTGGACCGGACCTGCACTGGGCTCGCAGCTTCTTTCACCTGTTTCCTG
AATTTTGTAAGAATATGTTCAGTCAATCCGTTTGCTGATCTTGTCCCATCCAACACAAGA
CGTCTTGCTGCCTCGAACGCAGGCCCATTGTAATGAGGAGCAAGTGGCCTGTATGCCTTG
TCTTGAGCATTCTGCTTGTCAACAATCAAAGCCATCTTCCGCAACGCTGCTTCAAACTGC
TCTTCTGAGATGATTCCATGGCTCAACCAATTCGCGCACAGCTGCGAAGAAATACGCAAA
GTAGCACGATCTTCCATGAGCTGCACGTCGCTGAGGTCAGGAACTTTAGAACAACCGACA
CCTTGATCCACCCACCGGACCACATATCCGAGCATGGACTGGCAATTCTCATCTAACTCC
TGCTCAATTTCAGCCGTACTCAACTGATTCTTCTCGAGCATGATTGGAGGGCTGAGAATT
TTCTTAAGCAATTCTTCGCGTGGCTGTAGTTTTAACGATTTT
>CL125837Contig1
GCGCGATCTCGTCCAGGAGGCCGTTGGCGAACTCGATGACCGCCTCGCCGCGGACGGGGT
TGTAGCCGGACGGCTGGGGGGCGGCGGCCTCCGACTTCCCCGAGAGGTCGAAGCCGGCCC
CGTCGGCTCCGACCTGCACCGCGGTGGACGGCACCACGTCGAAGCCGTAGAGCGCGTCGA
AGAGCGAGCCCCAGCGCGCGTTGACGGCGTTGAGCACGTAGCGCGCGTTGTCCGACGGGA
CGACGAGCTGGGGCGCCGGGATCTCCGCGATCTCGGGGTCGACGAACTGCG
>c516742_g1_i1
GGTGATACGTGCAAACATGGCCGCAGCAAATGGAATGCAACCCAAGCAGGTGTTGAGAGA
TGCGATCCGAGAGATTTGCATTGGATTGGAACCGAACAAAGAGGATGCATTTTGGCAAAG
CATGGAGGCGATGTTGCGGAAACACGTGGCCAGAAATGCTGAATTGTTGCGCATACGGGA
TGACATGCAGATGAAGATTGACGAATGGCACCAGAAACATCCGACCGTCACAGCAGAGAG
ATTAGGCGACAACAAGGATTACACAAATTTCTTAAAGCAAATTGGGTATCTTGTGCCGGA
GGAAACGGTACAGGCG
>c821725_g1_i1
GTCGGGATCCCGAAAAGGCAAGAGACTACTTGGCCGCCCGAATGGTCGGCCAAGCCTGGT
CGTGGCGGGCGAGGTTGCCGCTGACCAACTAGAGAGAGATACAACAAAGATGACTGTACA
ACTCGCTCACGTGAACGGGGAACAAAAAACATCGAGGTCGATCGTTCATGCACCAGGAGA
CAAAATCGACACAGAGCTGCGAGGCGAGGGACGGAGCAACAGGTGCCGGGAGAACTCCTC
GGCATCTTCGTGCCAACTCAGTCGGTGTGTCGAGAGTGCACGAACAGTCAACCTCGAACC
CTCAGTGCCAACGGCAGCCAACGGCAGCCAACGAGAGCCAACGAGAGCCAACGAGAGCCA
ACGAGAGCCTCAGAGCCTGGACTGCACCGCAGCCTTGGCAGCACGTCGTGCCTCCGTCAG
GGTGAACTCGGTGTAGCCATTGGGCACCTGCTTGCCGTTGAAGACGAGCTTCAAAGCAGC
CTGGAAGGCAATGTTGCTGTCGGTGTTGCCGGACATGGGCCTATACGCCTTGTCGGAGGA
GTTCTGCTGGTCGACGATCTTGGCCATCTCCACAAATGCATCCTTGATCTGGCTCTCTGT
GAGGAGGCCGTGCTGGAGCCAGTTGCCCAACAACTGAGAGCTGATTCGGAGGGTAGCTCG
GTCCTCCATCAGACCCACATTCTCCAGGTCGGGGACCTTCGAGCAGCCGACGCCCATGTC
GATCCAGCGCACCACGTAGCCCAAAATGGACTGAGCACTCTCGCGCAGCTCGTCAGTGAT
CTCCTTCTGGCTGGGCTTGGCCTTGAGCAACGGAGGCTCGAGGAGCATCTCGATCTTGGC

AGGTGGGCGCATCGCAAGCTGCATTTGGCGCTTCATGACGTTCACGCGGTGATAGTGGAT
GCTGTGGAGGGCACCGGCAGACGGAGATGGCACCCAGGCAGTCGTGGCGCCTGCCATCGG
TTCAGCGATCTTCGCCTCCAGCATTGCCTTCATGGAGTCGGGCTTGGCCCACATGCCCTT
GCCGATCTGAGCTTTGCCTTGAAAGCCACAGAAGAGGCCGATGTCCACATTCGCGGCTTC
GTAGGCCTTGATCCAGTCCTGCTGCTTCATGGCACCCTTGCGCACCACTGGGCCTGCCAA
GAAGCAGGTGTGGATCTCATCGCCAGTTCGGTCCAAGAAACCGGTGTTGATGAAGAACAC
ACGGTCGCTGGCAGCACGGATGCACTCGCGGAGGTTGGCGCTGGTTCGGCG
>c983328_g1_i1
GGTCTGAGCTGTTCTGTTCGTCAACCACTTTCGCCATCTTCGCAAAGGCTGATCTCACTT
GCTCTTCACTAACAACTCCATGCGTGAGCCAGTTTGCAAGGTACTGAGCGGAGATGCGAA
GAGTGGCACGATCTTCCATGAGGCCTGTTCCATTGATATCTGGAACTTTGGAGCAGCCGA
TACCTCGATCGATCCAGCGCACCATGTAT
>c1061531_g1_i1
GTGCTTTGGAGCTGACGCGGACGAACGTCTTGTCTTCATTGAGGGAGCGGGTCGCCTGCT
TGCCTCCCTTGTTGAAGGCGGTGGACAGCGTGCCTTGCATGAGCCCAGCCCAGTTCCTAT
ACACAGCGGCTTTGTCGCTCGCATCGACTGCGGCCACACTGTCCTCGCAATCAGCGATGG
CGGTAACGGCGGATTCAAGCAACACATCTTTGAC
>c783739_g1_i1
GCCTCCTTGCTCGGTGCCTGTCCGACCAAGGGTGGTGTCAAAAGATGCTCGAGGTTGCGG
GGCGGCCGCATGGCCAGCTGCATCTGCCTGGTCGGCACATCAACACGGTGGTAGTGGATG
CTGTGCAACGCCGCCGCAGATGGCGAGGGCACCCAAGCAGTGCTCGCACCGGCCATCGGC
TCAGCGATCTTGGCCTCGAGCATGCCCTTCATGCTGTCTGGCTTGGCCCACATGCCCTTG
CCG
>c190916_g1_i1
ATAGTCGTTACAGCAGCTTCAAGAACTAAATCTTTTACGCCAGCGGCATCTGTTTTACCA
ATTGGGCTGTGCGCATCAATTTCAATTACAACGTGTAGGCCGTTGTTTAAAAGTACGATT
TCTGTTGGAGCCGATTCTTCGCCATTAAAACCAACAAATTGTGCTTCTTTAGCCAAAGTC
GATGTCGAACCGTCTTTTAACGT
>c603031_g1_i2
GCACCCCCCGCGTGAGGCGCTCCTTGCAGCCGTCCTCTGCCCGCCTGTGATGTCCAAGGC
CGAGCGGCCGTCCGCGGCAGAAGTCCAGCAGGAGCTGGAGGAGAGCTGCCAGTCGATCCT
GGGCTACGTGGCCCGCTGGGTAGAGCAGGGCGTCGGCTGCTCGAAGGTGCCGGACCTTCA
GGGCGTCCAGCTCATGGAGGACCGCGCCACCTTGCGCATCTCGTCGCAGCTGCTGGCCAG
CTGGCTGCGCCACGGCGTCGTCACCAGCGAGCAGCTCGACGCCGCCATGCTCAAGATGGC
CCATGTGGTCGACGGACAGAACGCCAACGACGCCGCCTACCGCCCCATGTCCACG
>c775803_g1_i1
GTCCAGGATGGTGGTGAGGGGACCCTCGATAGTGATGTCCTTGATGCCGCTCAGGGCTGC
CGCACCGATCGGGTCGGCGCGGTCGACCTCAAGGATCAGGTGCATGCTGTTGTGCTTGAG
GAAGATGCGGCCCTGGTCAGGGACGGCCTGAGTGGGGTCAGCACCCTGGGGCTTGGCCGC
AGGTGGGCCCATGCTGCCGACGGAGCCAACGAAGAGAGCCGGAGTGGCCAGGCTGGTGGT
CTCGCCGCTCTTGAGCTGCAGCTCCAACTGCTGGGAGGCACCCACAAACTTGGGCCAGAG
GCGAGTCACCTCGGACCATTTGCCCGTGTTCAGAGGGGCGATTTCGTCGAGAATGCTGTT
GGCGAACTGGCGCACGGCCTCACCTCGGACAGGGTTGTAGCCGCCGGCCTTGGGGGTGCT
CTCGCCAACCGGCTGCATCGACGCCTGGGGGACGACGTCGAATCCGTACAGAGCATCAAA
GAGAGAGCCCCAGCGAGAGTTGAGGGCGTTCAGGACGTATCGTGCATTGTCGACCGGCAC
GACGAGCTGAGGAGCAGGAATGTCTGCAATTTCGGCGTCGACATTCTGGGTGCTGATGGT
GAGCGTTCCCTTGTCCTCCTCAAGGTAGCCAATCTCTCGGAGAAACTTGTGGCACATGGG
GCGAGTGGCTGGGGATGTGGGGTCAGTGCCCTCGGATCGCACCTTTTGGTAGAAGTTGTC
GATCTTTACCTGCAGATCGTCACGCTGCTTCAGGCAAGACTCAAGTTCAGGCTGGAGGTC
CTTGATCAACTTGTTGAGGCTGCCCCAGAAGTATTTGCTGCTGAAGCCGGTGCCAGGGCA
CAACTTCTCCTCCACGAGTTTAGCCAGGGTCGGATGGATCTTGAGGCCACCGGACTCCAC
GTAGTTGGAGGGCACTGGGACGACTGCGGACATTGTGGTCAACTCGCGCACTGAGTGTCA
GGGTGTGACAGATGATGGATGATGGGGGTGGCCTTGAGT
>c527602_g2_i1

GAAGAATTCTTTGGACTCCGTCGAAATACCATCAAGATGGGAGTGATGGACGAGGAGCGC
AGGACCAGCGTTAATCTTCGGGAGTGTCTTCGAGTGGCGTCTGAGCGTGTGTTCTTTATC
AATACTGGCTTCCTTGATCGTACCGGTGACGAGATCCACACTTGCATGCATGCAGGACCT
GTTGTTCCTAAAGCTGCAATGAGGCAACAAATTTGGATCAAAGCCTACGAAACGAGCAAC
GTTGACGTTGGTATTGCAGCTGGAATGTTGGGCCGAGGCCAAATTGGCAAAGGAATGTGG
GCCAAACCTGACAGTATGAAAGCAATGTTGGAAGCGAAGATAAGCGAGCTCATGGCGGGA
GCAAGTTGTGCTTGGGTTCCATCACCGACAGCGGCAACTTTGCATGGAATTCATTACCAC
CGCGTTGATGTCTTGGCAAGGCAGGGGCAGCTGAGCACGCGAGAGCCAGCCAAGGTTGAA
GAGATATTGCAGCCTCCTCTGTTGAAGGAGTCCTTGAGCAGGGAAGCGATTGAACACGAG
TTAAAAGAAAACGTGCAAAGTATATTGGGATATGTCGTGCGTTGGGTCGATCTTGGTATC
GGTTGCTCGAAAGTTCCTGACATGTCGAATGTGGGATTGATGGAAGACAGAGCAACTCTG
AGGATATCATCACAATTGTTGGCGAATTGGTTGGCTCATGGATTGATTTCTGCTGAAGAA
CTTAAGCAAACCTTTAAGGAGATGGCGGTACTTGTAGACCAGCAGAATGCGCAGGACAAG
GCTTACAAACCCATGATTACAAACCTCGAAGAAAATATTGCATACAACGCTGCAATGACT
CTCGTATTGCAGGGTCACACAACACCAAACGGCTACACTGAGCCCATTCTACATGAAGCC
AGGCGAAAGGTCAAGGCTGCAAGCATGTCGTCTAGATTGTGATCAAATGGACATAAGTTC
CAAGCTTTAATGGTTAGAAG
>CL229Contig8
CTGCCATGCTACACCAGCGCCTGCAGACCGAAATCTCCGCATGCAGCGGTGCAAATCAAA
AGAAAGAAAAAGAGAGAGTGCTCTCAGAGTCTGGAAAGCCGTCCAAGCTGAGCCTTGGCG
GCACGCCTGGCCTCATGTAGCGTGTACTCCGTGTAGCCATTTGGCGCTCTCTGGCCCTCC
AGCACGAGCTTCAAGGCTGCATTGAAAGCAATGTTATTATCTGGGTCTGTAGACATGGCT
CTGTAACCTCGGTCTCGAACGTTTTGTTGGTCCACGACCTTAGCCATTTCCAGAAATGTC
TTCCGCAACTGGGGCTCGGTGACAAGACCGTGACGCAGCCAATTCGCCATAAGCTGCGAG
GAGATTCTCAGTGTTGCTCGGTCCTCCATGAGACCCACGTTGGAAAGGTCAGGCACCTTT
GAACAGCCCACGCCAAGGTCTACCCAACGCACCACGTAGCCTAGGATGCTCTGGGCATTT
TCCCGAAGCTCGTGCTCGATCTCCTGTTGCGAAGGCTGGCTGCTGAGCAGCGGTGGTTCC
AGGAGCAGTTCCATCTTCGCTGCGGGTCGCATGGCCAACTGCATCTGTCGCTCTTGCACG
TTCACGCGGTGATAATGGATGCTGTGGAGGGCTGCAGCCGACGGGGAAGGAACCCAAGCT
GTGCTCGCTCCAGCCATGAGCTCTGAAATCTTGCCTTCCAGCATAGCCTTCATGCTGTCT
GGCTTTGCCCACATGCCTTTTCCAATCTGCCCCTTCCCCTGGAAACCAGAGTGCAAGCCA
ATGTCGACGTTGCCGGCCTCATAGGACTGGATCCAGGGTTGTTTTTCATGTCAGCCTTC
ATGACGACAGGACCTGCGGCCATGCATGTATGTATTTCATCGCCGGTGCGGTCCAAGAAG
CCAGTGTTGATAAAAAAGACCCTTTCTGCTGCTGCGCGAATACACTCCCTGAGGTTTGCA
CTGGTGCGGCGTTCCTCGTCCATGATGCCCATCTTGATTGTGTTCCTCCTCAGGCCGAAG
AATTCCTCAGCCCGTCCAAAGAGCCGGCCTGTGAGTGCAATCTCCTGGGGACCATGCTGC
TTCGGCTTGACTATATAAAGCGAACCAGCGCGGCTATTCATCACTCGGGCTGTGCCTCGT
AGGTCAGGCAGGGCTGCAGCTGCAGTGACGAGGCAGTCTACGAATCCTTCAGGGAGTTGT
TTCCCGCTAGAGGTGAGCACCATGTCAGTAAACATGTGGTGCCCCACATTGCGCACCAAC
GCCACCGCTCTGCCAGGAAGGGTGTTGGCCTTGCCCTGGGCATCGCGAAAGGCGACATCT
GCGTTGAGGCTCCTGGTCATAGCTTTACCATCCTTGACAAATGGAGCTTCCAGGGTTCCA
CGAAAAACACCACAAATATTGCCGTAAACTCTTGATTTGTCTGCAGCATCCACTGCTGAC
ACTGAATCCTCCATGTCAAGGATAGTGGACAGCGCCGCCTCAACTGTGATGTCCTTGATG
CCGCTCAAGGCGCTCTGGCCGATCGGATCGCTACGGTCAATTTCTAAAATGAGATGCAAG
CCATTGTGTTTCAGGAATACTCGACCAACATCTGGGACACTTGGACTCGCAGCTTGGGGA
GCCACGTCAGCATCTGGAGGCCCAAGGTTGCCAGTGGAGCCCACAAACAGTGAGGGAGTC
CTAAGACTCGTGGTGGCACCACTTCTTGTGAGCAGTTCCAAGCGCTGCTCGGTGCCCACA
TACTTGGGCCAGAGCCGGCAGACATCAGCCCAGGCCCCAGACACCAACGGTGCGATTTCG
TCTAGCAGGCCATTTGCAAACTCAATCACAGCCTCTCCACGCAATGG
>c974883_g1_i1
GGTAGAACCGGACTCCATGGCTCGTATGTTGAAGGAGAAGCACAACCACCCATCTATGGG
TGCCACCACTGCATGGGTGCCGTCTCCCACTGCCGCAACACTACATGCTCTCCACTACCA
CCAGGTCGACGTCTTCGCTGTGCAGAAAAAGCTTGCTGCCGGTGGTCCACGTGCCTCTGT
GGATGCCATCCTCACACCTCCAATGCTAGAATCCACACTGAGTGCAAGCGAGATCACCTC

TGAGCTCGAGACCAACGCACAGGCTATTTTGGGTTACGTTGTCCGATGGGTGGAGCAAG
>c666275_g1_i1
CCGGCTTGGCCCACATGCCCTTGCCAATTTGACCCTTTCCTTGAAAGCCTGAGTGAAGGC
CGAGGTCTACGTTGCCAGCTTCGTAGGACTTGATCCACGGCTGCTTCTTCATGTCAGCCT
TCTTCACTACAGGACCTGCCAACATGCACGTGTGGATCTCGTCACCAGTGCGGTCCAGGA
AGCCTGTGTTGATGAAGAAGACCCGCT
>c528479_g1_i1
GTGAATCCTTAAAATAATAACACATGGCACTGCGCACACCAACACATGCAACAAGATATG
CGCAACAACACATGGCACCGTACACACCAACACATGCAACAAGGGATGTGGGCAAAGCCA
GACGAAATGGCAGCCATGATAGCATCCAAAGGTTCTCACCCGCAATCGGGCGCGACCACG
GCCTGGGTCCCCTCACCAACTGCTGCTACTCTTCATGCGATTCATTATCACGAGACAAAT
GTGCTCAAAACGCAAAAGCGCATGCTTGCTGAGGGACGGCGTGCCAAGCTGCAGGACATT
CTATTGCCTCCATTGATGAGTCTCAGCTCCGCAGGCCCTGTAGCACTCACTCAAGATGAG
ATTCGTAAGGAGATGGAAAACAAC
>c927326_g1_i1
GTCCATTGTGCTTGATGAAGATGCGGCCCTGATCTGGCATGGCCTCTGTGGCGGCCGTGC
CCTCCGGCTTGGCTGCCGGTGGACCCATGCTGCCGACGGACCCGACGAAGAGGGAGGGGG
TGGCCAAGCTGGTGCTCTCGCCGCTCTTCAGCTGGATCTCCAACTGCTGCGAGGCACCGA
CGAACTTGGGCCAGAGCCGTGTCACCTCGGACCACTTGCCGACATTGAGAGGGGCGATCT
CATCCAACAAGCCGTTGGCGAACTGGCGCACAGCTTCGCCTCTCACGGGGTTGTAGCCAC
TGGTGGCGTGAGGGGTGCTTTCGCCGACAGGCTGCACCGCAGACTGCGGGATGACATCGA
ATCCGTATATGGCGTCGAAGAGCGAACCCCATCTTGAATTGAGAGCATTCAACGCGTAGC
GAGCGTTGTCCACAGGCACGACGAGCTGGGGGGCAGGGATGTCGGCGATCTCCGCATCGA
CATTCTGAGTGCTAATGGTGAGCGATCCTTGATCCTCCTCCAGGTACCCGATCTCCCTCA
GGAACTTGTGGCACATCGGGCGTGTGGCAGGAGAGGCAGGGTCGGTTCCCTCGGACTTGA
CCTTCTTGTAGAAGTCATCGATTTTGGCCTGCAGGTCGTCCCGGTGCTTCAGGCAAGACT
CCAGCTCAGGTCCGAGCTCCTTCACCACCTTCTCCAAACAGTTCCAGAAGTACGTGCTGC
TGAACCCAGTACCAGGGCAGACCGCTTCCTCAACCAGTTTCGCCAATGTCGGATGGACCT
TCATTCCACCAACCTCGATGTAGTTCGAAGGAACATTCACGACGGTCGACATGGTTGAAG
TTTGGGCGGCAAACAGGGCCGAAAGGTGTGGGACTTGGGGGAATG
>CL5221Contig1
CGCCTTGCGCCGGAACGCGTGCAGCACGGGCTCGGTGAGGCCGCTGGGCAGCCGGCAGCC
ATCGCGCACGAGCCGACCGGCCGCCTGGAATGCCAGGGACTGGTCCAGAGCCGCGCACAT
AGGACGATAGGCGGCGTCGCCAGCGTTCTGCTCGTCGACCACCCGGGCCATCTTCCGCAT
GGCCGCATCGAGCTGCGCGTCGGTGACGACACCGTGGCGCAGCCAGCTGGCCAGCAGCTG
CGACGAGATGCGCAAGGTGGCGCGGTCCTCCATGAGCTGGACGCCCTGAAGGTCCGGCAC
CTTCGAGCAGCCGACGCCCTGCTCTACCCAGCGGGCCACGTAGCCCAGGATCGACTGGCA
GCTCTCCTCCAGCTCCTGCTGGACTTCTGCCGCGGACGGCCGCTCGGCCTTGGACATCAC
AGGCGGGCAGAGGACGGCTGCAAGGAGCGCCTCACGCGGGGGGTGC

**synthase**

| ID | Annotation |
|---|---|
| CL48332Contig1 | isocitrate lyase |
| CL67599Contig1 | isocitrate lyase |
| CL125104Contig1 | isocitrate lyase |
| CL125491Contig1 | isocitrate lyase |
| CL1Contig2463 | isocitrate lyase |
| c664616_g1_i1 | isocitrate lyase |
| c892671_g1_i2 | isocitrate lyase |
| c914354_g3_i1 | isocitrate lyase |
| c914354_g3_i2 | isocitrate lyase |
| c914354_g3_i3 | isocitrate lyase |
| c909308_g1_i1 | isocitrate lyase |
| c692422_g1_i1 | isocitrate lyase |
| CL1Contig10513 | isocitrate lyase |
| CL1Contig12374 | isocitrate lyase |
| c775297_g1_i1 | isocitrate lyase |
| c883116_g1_i1 | isocitrate lyase |
| c849717_g1_i1 | isocitrate lyase |
| c902440_g2_i1 | isocitrate lyase |
| c738735_g1_i1 | isocitrate lyase |
| c810828_g1_i1 | isocitrate lyase |
| c892052_g1_i1 | isocitrate lyase |
| c810024_g1_i1 | isocitrate lyase |
| c398051_g1_i1 | isocitrate lyase |
| c810371_g3_i4 | isocitrate lyase |
| c524822_g1_i1 | isocitrate lyase |
| c736104_g1_i1 | isocitrate lyase |
| c836096_g1_i1 | isocitrate lyase |
| c736768_g2_i1 | isocitrate lyase |
| c892052_g1_i2 | isocitrate lyase |
| c907594_g1_i3 | isocitrate lyase |
| c357732_g1_i1 | isocitrate lyase |
| c933242_g1_i1 | isocitrate lyase |
| c905333_g1_i2 | isocitrate lyase |
| c831242_g1_i1 | isocitrate lyase |
| CL12084Contig2 | isocitrate lyase |
| CL53156Contig1 | malate synthase |
