## Supplementary material for "Caloric Restriction Remodels Energy Metabolic Pathways of Gut Microbiota and Promotes Host Autophagy": SI-TableS4

**Supplementary Table S4. The primers' list for RT-PCR reaction**

| Gene symbol | Oligo sequence (5'→3') |
|---|---|
| MDH1-F | GTTCATCACGACTGTGCAGC |
| MDH1-R | GATTACGCCCATCGACACGA |
| MDH2-F | CCCAAGGTTGACTTTCCCCA |
| MDH2-R | TAAGCCATGGACAGAGTGGC |
| Me1-F | ACCCGCATCTCAACAAGGACT |
| Me1-R | ACAGAAGATACCTGTCGAAGTCA |
| Idh1-F | CGGGTGTGAGCGGGGTTAT |
| Idh1-R | ATTGGTGGCATCCCGATTCT |
| IDH2-F | CGGAGCTATGACCAAGGACC |
| IDH2-R | CTACAACGCCCACCCTCTAC |
| GAPDH-F | TCTCTGCTCCTCCCTGTTCT |
| GAPDH-R | TACGGCCAAATCCGTTCACA |
| Pdk4-1F | GCAGTCCTCACCAACCCTAC |
| Pdk4-1R | CTGTAGCGGGAGAACAGCTC |
| PDC4-F | AGCATCTGCCCTCTGTTGAC |
| PDC4-R | TTACAGCGTCAGCGATCTCC |
| Pdk2-F | CACCCCAAACACATTGGCAG |
| Pdk2-R | AGGGGACGTAGACCATGTGA |
| PC-2F | TGGAGACTGTGGTGACTTCG |
| PC-2R | GAGGGCCATTGCAGGTAGTG |
